# scDIVA: semi-supervised integration and fine-grained annotation of tumor-immune single-cell atlases

**DOI:** 10.64898/2026.08.07.743520

**Authors:** Viraj Rapolu, Alireza Karbalayghareh, Brennan Lee, Wilfred Wong, Christina S. Leslie

## Abstract

Single-cell atlases of the tumor-immune microenvironment have defined numerous fine-grained immune cell states, but each study uses its own nomenclature and procedure for annotating cell types. Transferring annotations from a reference atlas to a query dataset is complicated by both batch effects and by the presence of query populations that the reference does not contain. Here we present scDIVA, a semi-supervised deep generative model that adapts the Domain Invariant Variational Autoencoder to scRNA-seq for fine-grained tumor-immune label transfer. Three encoders disentangle each cell’s expression profile into separate latent subspaces for cell type, batch, and residual variation; a single decoder reconstructs the cell’s expression profile from all three latent embeddings, and auxiliary classifiers on the cell type and batch embeddings encourage each encoder to capture the respective source of variation; scDIVA’s cell type embeddings are batch-invariant by construction rather than through explicit or adversarial correction. Benchmarked against four established reference-mapping approaches—Harmony/Symphony, scANVI with scArches, scPoli with scArches, and Seurat with label transfer—across six tumor-immune atlases spanning five cancer types, scDIVA achieved the highest mean macro-F1 in five of six atlases and the highest biological conservation scores. To detect query-enriched populations, we adapted the Milo differential abundance (DA) framework, added a directional test, corrected spatialFDR weighting, and parallelized neighborhood distances for atlas-scale data, and applied it to scDIVA’s cell type embeddings with reference-versus-query membership as the condition. This design correctly identified cell types held out from the reference as OOR and flagged exhausted CD8 T cells from a tumor-immune atlas as OOR relative to a healthy pan-tissue immune reference; conversely, this procedure confirmed a conserved immune landscape between two independent colorectal cancer cohorts. scDIVA thus couples fine-grained annotation and integration with an FDR-controlled test for states the reference lacks, solving both tasks required for accurate label transfer in tumor-immune scRNA-seq atlases.

## Introduction

Single-cell RNA sequencing (scRNA-seq) has transformed our ability to dissect the cellular composition of complex tissues and tumors, in particular enabling the discovery and characterization of novel immune cell types and states, both in health and in disease^1–3^. Within the T cell compartment alone, single-cell analysis has defined diverse phenotypic states—CD4+ and CD8+ naive, effector, and memory cells; CD8+ progenitor and terminally exhausted cells; distinct classes of regulatory T cells; and proliferating subsets of several of these populations—with distinct functional programs and specialized roles in antitumor immunity, autoimmune pathology, and response to immunotherapy^4–8^. Large-scale profiling efforts have generated scRNA-seq atlases of immune cells in healthy human tissue, notably the CellTypist resource generated through the Human Cell Atlas^2,9^, representing a major step towards building a comprehensive catalogue of healthy immune cell types. However, a systematic, fine-grained, and consistent annotation of immune cell states in disease, particularly in cancer, is not yet available. While numerous scRNA-seq atlases have profiled the tumor immune microenvironment (TME), each study tends to use its own nomenclature and preferred marker genes, making the mapping between cell types in different datasets challenging^10^.

Transferring fine-grained immune cell annotations from a reference atlas to an independent query dataset requires solving two problems. First, datasets generated across laboratories, platforms, and tissue-processing protocols contain systematic batch effects that obscure biological signal and must be accounted for before any cross-dataset comparison or integration can be performed^11^. Second, a query dataset may contain populations that the reference lacks, and unless these out-of-reference (OOR) populations are detected, they may be assigned an inappropriate reference label; systematic benchmarking of tumor immune atlases has shown that missing reference subtypes, including exhausted and memory T cell states, are a recurrent source of cross-mapping error^10^. These problems are coupled: detection of OOR populations in a separate dataset requires batch correction, as it may be unclear if such populations are genuinely novel or differ from the reference due to technical artifacts.

While numerous existing approaches address aspects of the integration and annotation problem—batch correction, label transfer, and various deep learning embedding algorithms—obtaining accurate and fine-grained annotations on a new dataset still often requires extensive manual checking and trial-and-error. Unsupervised methods such as Harmony^12^ produce batch-corrected embeddings but require a separate cell type classification procedure and lack any mechanism for detecting OOR query cells. Symphony^13^ provides a compressed representation of a Harmony-corrected reference dataset and enables mapping of query cells to this embedding for transfer of reference-defined annotations but does not perform OOR population detection. Seurat provides anchor-based integration and label transfer within a single framework^14^ but similarly has no formal mechanism for flagging OOR populations.

Deep generative models were first introduced in single-cell transcriptomics via scVI^15^, a variational autoencoder that embeds cells while accounting for batch effects through a batch covariate in its generative model. scANVI^16^, a semi-supervised model, extends this framework by joint integration and classification of cells in a shared latent space, transferring cell type labels to unlabeled query cells. The scArches^17^ transfer-learning framework adapts these models for reference mapping onto an independent query dataset, while scPoli^18^ adds batch-aware prototype representations and treeArches^19^ includes a hierarchical classifier with cell-level rejection. These related methods implicitly handle batch effects by conditioning the decoder on a batch covariate rather than separating batch-related from cell-type-related expression variation in the model. A parallel line of work makes this separation explicit. For example, scDREAMER^20^ and biolord^21^ remove batch effects adversarially or at the attribute level, while DRVI^22^ learns disentangled latent factors without supervision. Another model, inVAE^23^, partitions the latent into invariant biological and spurious technical representations and conditions its priors on biological covariates. Finally, a recent preprint presents FADVI^24^, which uses dedicated batch, label, and residual latent subspaces, placing standard Gaussian priors on all three and enforcing their separation through supervised classification heads, adversarial heads with gradient reversal, and a cross-covariance penalty.

To detect query populations that the reference lacks, various methods assess the OOR status at the cell or population level. At the level of individual cells, classifiers may report confidence scores or perform explicit rejection. Prior strategies include softmax probabilities over the reference labels, prototype-based uncertainty in scPoli, and cell-level rejection in treeArches^18,19^. These methods assess which reference type a query cell most resembles and then whether to support non-assignment. However, a genuinely novel query cell could still receive a high score for a related reference type, and information is not shared across individual query cells. Dann et al. first developed Milo^26^, a differential abundance (DA) testing procedure that tests for neighborhoods enriched in one condition relative to another, and subsequently applied this framework to human populations to detect disease vs healthy^25^. We leverage this strategy to identify query-enriched neighborhoods as candidate OOR states, using the fraction of query cells per neighborhood to distinguish truly unseen states from populations merely enriched in the query dataset. Another method, contrastiveVI^27^, isolates target-specific (disease) variation in a dedicated salient latent space, which it analyzes by clustering and differential expression rather than DA.

More recently, single-cell foundation models pretrained on extensive scRNA-seq corpora have emerged to learn general-purpose, transferable cell representations applicable across tissues and conditions. Most are trained by self-supervision, learning to predict held-out gene or expression tokens without using cell-type labels: scBERT^28^ and Geneformer^29^ mask and reconstruct discretized expression values or gene rankings, scGPT^30^ applies a generative objective over binned expression, and scFoundation^31^ models continuous counts with a masked autoencoder. The Universal Cell Embedding^32^ extends self-supervised masked prediction across species using protein-language-model gene representations. Meanwhile, pretraining corpora for such single-cell foundation models have grown from roughly one million cells to over one hundred million^33^. Annotation is performed afterwards rather than during this label-free pretraining, either by fine-tuning a supervised classifier on labeled cells or by transferring labels from an annotated reference using the pretrained embedding. A related approach, SCimilarity^34^, instead learns its embedding through supervised metric learning over annotated cell types, enabling large-scale search for transcriptionally similar cells alongside dataset integration. These foundation models are designed for broad label transfer rather than fine-grained reference-to-query annotation. None separate technical from biological variation into dedicated subspaces, and none provide a population-level test to detect cell states absent from the reference. Systematic evaluations report that single-cell foundation models plateau in performance well below current corpus sizes, do not exhibit the data-scaling behavior of large language models, and do not consistently surpass task-specific baselines on cell type annotation and integration^35^. In zero-shot pipelines, the embeddings of leading models have been found to fall below standard pipelines such as highly variable gene (HVG) selection with Harmony or scVI^36,37^.

Here, we present scDIVA, a semi-supervised deep generative model that addresses integration, classification, and OOR cell state detection in a unified framework. scDIVA builds upon prior work that explicitly attempts to disentangle biological from technical sources of variation but takes a semi-supervised strategy to partition the latent space. Specifically, scDIVA adapts the Domain Invariant Variational Autoencoder (DIVA), originally developed for domain generalization in image classification, to scRNA-seq data^38^. Three encoders disentangle sources of single-cell gene expression via distinct latent subspaces for cell type (*z_y_*), batch identity (*z_d_*), and residual variation (*z_x_*). A single decoder reconstructs the expression profile from the concatenated representations, and auxiliary classifiers on *z_y_* and *z_d_* encourage each encoder to capture the appropriate source of variation. Unlike scVI, scANVI, or scPoli, which condition on batch, or scDREAMER, which removes batch adversarially from a shared space, scDIVA gives batch-related variation a dedicated latent subspace. scDIVA trains end-to-end in a single pass without a pre-trained model, simplifying the training pipeline relative to scArches-based reference mapping. The recently preprinted FADVI^24^ similarly partitions the latent space but enforces subspace independence through adversarial and penalty terms on standard priors, whereas scDIVA conditions the cell type and batch priors on their respective labels and uses no adversarial or decorrelation terms.

The structural latent separation in scDIVA enables a principled approach to detecting cell states in the query that lack reference counterparts, as we apply a reference-versus-query DA framework from Dann et al.^26^ to scDIVA’s cell type embedding. By treating membership in the reference or query set as the experimental condition and performing DA analysis on the *z_y_* embedding, we identify neighborhoods that are statistically enriched in the query relative to the reference. Because batch variation has been structurally separated into *z_d_*, differential abundance in *z_y_* reflects genuine differences in cell state composition rather than technical artifact—a property that would not hold if cell type and batch information were entangled in a shared latent space. This approach produces continuous log-fold change estimates and formal p-values at the neighborhood level with FDR control, providing a statistical framework for evaluating potential OOR cell states.

We applied scDIVA to six pan-immune tumor atlases spanning five cancer types and comprising up to 412,000 cells per dataset. Through controlled holdout experiments and cross-context mapping from healthy to tumor-derived immune cells, we found that the DA framework identifies OOR cell states, including tumor-specific populations such as exhausted CD8+ T cells that have no counterpart in healthy immune cell references. Conversely, upon integration of two independent cohorts of the same tumor type, this framework reported a conserved immune landscape with few query-enriched neighborhoods, demonstrating that it flags OOR cell states only when the underlying biology differs. In systematic benchmarking against four established reference-mapping approaches across these atlases, scDIVA achieved the highest mean macro-F1 in five of the six atlases, significantly outperforming all four competitors in four of them and Harmony/Symphony in all six. On global composite metrics aggregated across datasets, scDIVA significantly outperformed Harmony, Seurat, scPoli + scArches, and scANVI + scArches on both biological conservation and batch correction. Together, these results establish scDIVA as a purpose-built tool for accurate fine-grained immune cell type annotation, integration, and cell state discovery in the tumor microenvironment.

## Results

### scDIVA disentangles batch and cell type into separate latent subspaces

scDIVA adapts the Domain Invariant Variational Autoencoder to scRNA-seq by mapping each cell’s expression profile through three encoders into three latent subspaces: one for cell type (*z_y_*), one for batch identity (*z_d_*), and one for residual variation (*z_x_*). A single decoder reconstructs the profile from the concatenated representations, and auxiliary classifiers on *z_y_* and *z_d_* drive each encoder to capture its assigned source of variation (**Fig. 1A**). The model assumes that batch and cell type are conditionally independent given a cell’s expression profile. In our experiments, we did not present scDIVA with held-out data after model training but instead used the model in a semi-supervised and transductive setting. Namely, scDIVA was trained in a single pass on the combined reference and query, where only reference cells carried cell type labels, while query cells were represented by their expression profile and batch identity without a cell type label. Query cell types were then annotated with the *z_y_* classifier.

**Figure 1.**
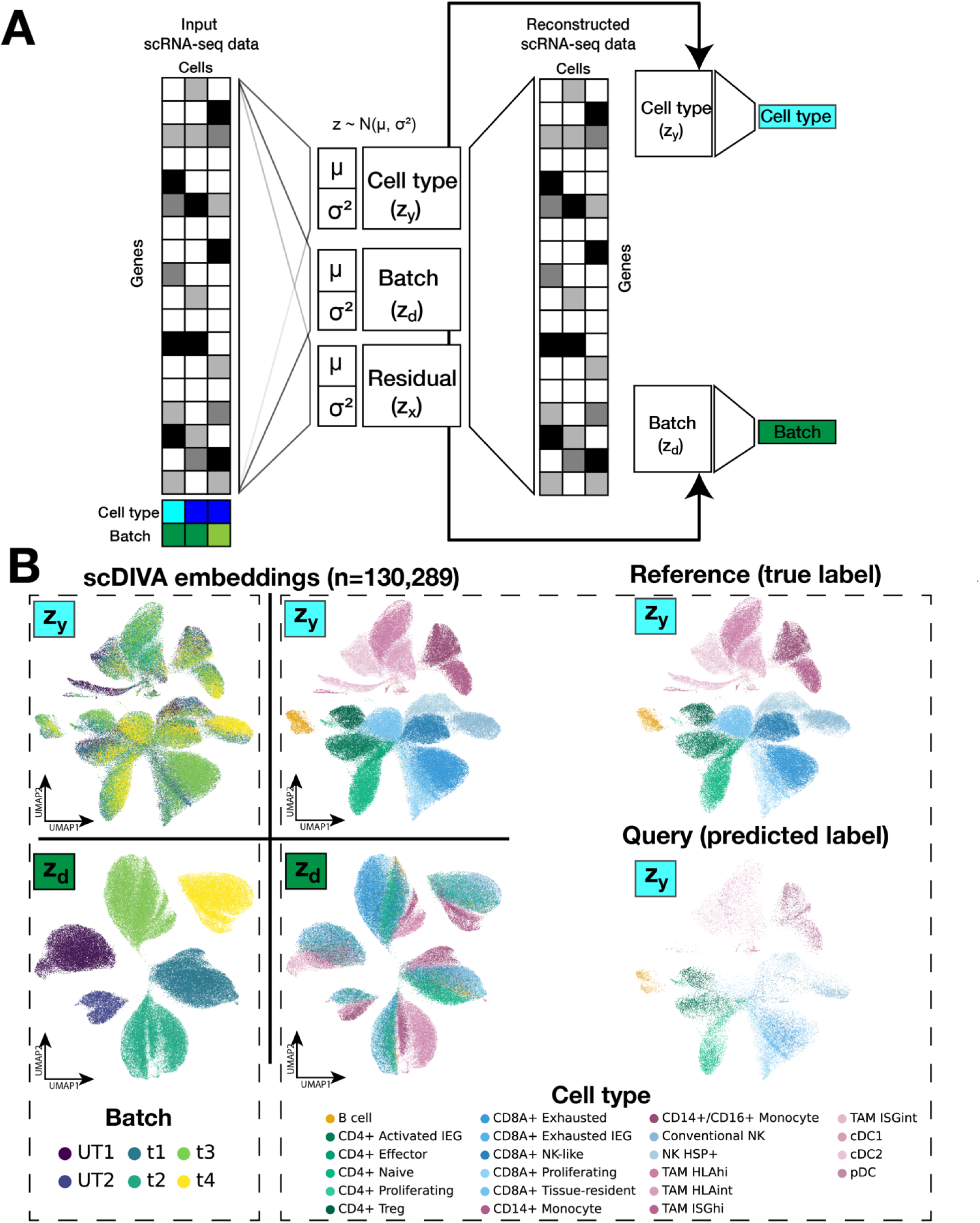
scDIVA disentangles batch and cell type into separate latent subspaces. (A) Schematic of the scDIVA architecture. Three separate encoders *q_фy_*(*z_y_*| x), q_фd_(*z_d_*| x), and q_x_(*z_x_*| x) encode cell type, batch, and residual variation from single-cell expression profiles. A use the latent space embeddings *z_y_* and *z_d_* to predict cell type and batch labels respectively. single decoder reconstructs the input from the latent space representations. Auxiliary classifiers (B) Left: UMAPs of scDIVA *z_y_* and *z_d_* embeddings for the Krishna/Hakimi ccRCC immune atlas, colored by batch (left) and by cell type (right). *z_y_* mixes cells across batches while separating cell types, whereas z**_d_** separates batches while collapsing cell types. Right: UMAPs of scDIVA *z_y_* embeddings colored by ground-truth annotation (top) and held-out query cells colored by scDIVA-predicted labels (bottom). As a semi-supervised model, scDIVA was trained on all six batches (patients), with one held-out batch as the unlabeled set or query set. Cells in the query set contributed gene expression profiles and batch identity but not cell type labels.

As this model explicitly disentangles batch and cell type, we demonstrated this disentanglement by training scDIVA on a clear cell renal cell carcinoma (ccRCC) tumor-immune atlas (Krishna/Hakimi^39^) that profiled multiregional samples from 6 patients, with each patient sample treated as a batch. We treated 5 patients as the labeled reference and one held-out patient as the unlabeled query and confirmed that the *z_y_* embedding mixes cells across batches while grouping cell types, whereas the *z_d_* embedding separates batches while mixing cell types (**Fig. 1B**). The residual space *z_x_* is structured by neither and captures the variation left over once batch and cell type are accounted for. The predicted labels for the held-out cells recovered the ground-truth structure in *z_y_* (**Fig. 1B**). Repeated across each of the six held-out batches, scDIVA achieved a mean overall accuracy of 0.842 and mean macro-F1 of 0.787. The confusion matrix of aggregate predictions across the six scDIVA runs on the Krishna/Hakimi atlas dataset displays a clean diagonal (**Fig. S1**).

### scDIVA improves fine-grained classification and integration across tumor-immune atlases

We benchmarked scDIVA against four published reference-based cell annotation approaches: Harmony with Symphony query mapping, scANVI with scArches, Seurat label transfer, and scPoli with scArches. Benchmarking was performed on six tumor-immune atlases spanning five cancer types and up to 412,000 cells—Pelka/Hacohen^40^ (colorectal cancer), Krishna/Hakimi^39^ (ccRCC), Chan/Rudin^41^ (small-cell lung cancer), Leader/Merad^42^ (non-small cell lung cancer), Vázquez-García/Shah^43^ (ovarian cancer), and Zhang/Yu^44^ (colorectal cancer)—using matched reference/query splits (**Fig. 2A**, **2B**). Because fine-grained immune labels are frequency-imbalanced, overall accuracy is dominated by a handful of abundant populations. We therefore computed macro-F1, the unweighted mean across cell types of each cell type’s F1 score, as the primary measure of annotation quality, since it weighs every cell type equally and penalizes both missed and over-called populations. We additionally report macro-recall as a supplementary evaluation metric (**Fig. S2**).

**Figure 2.**
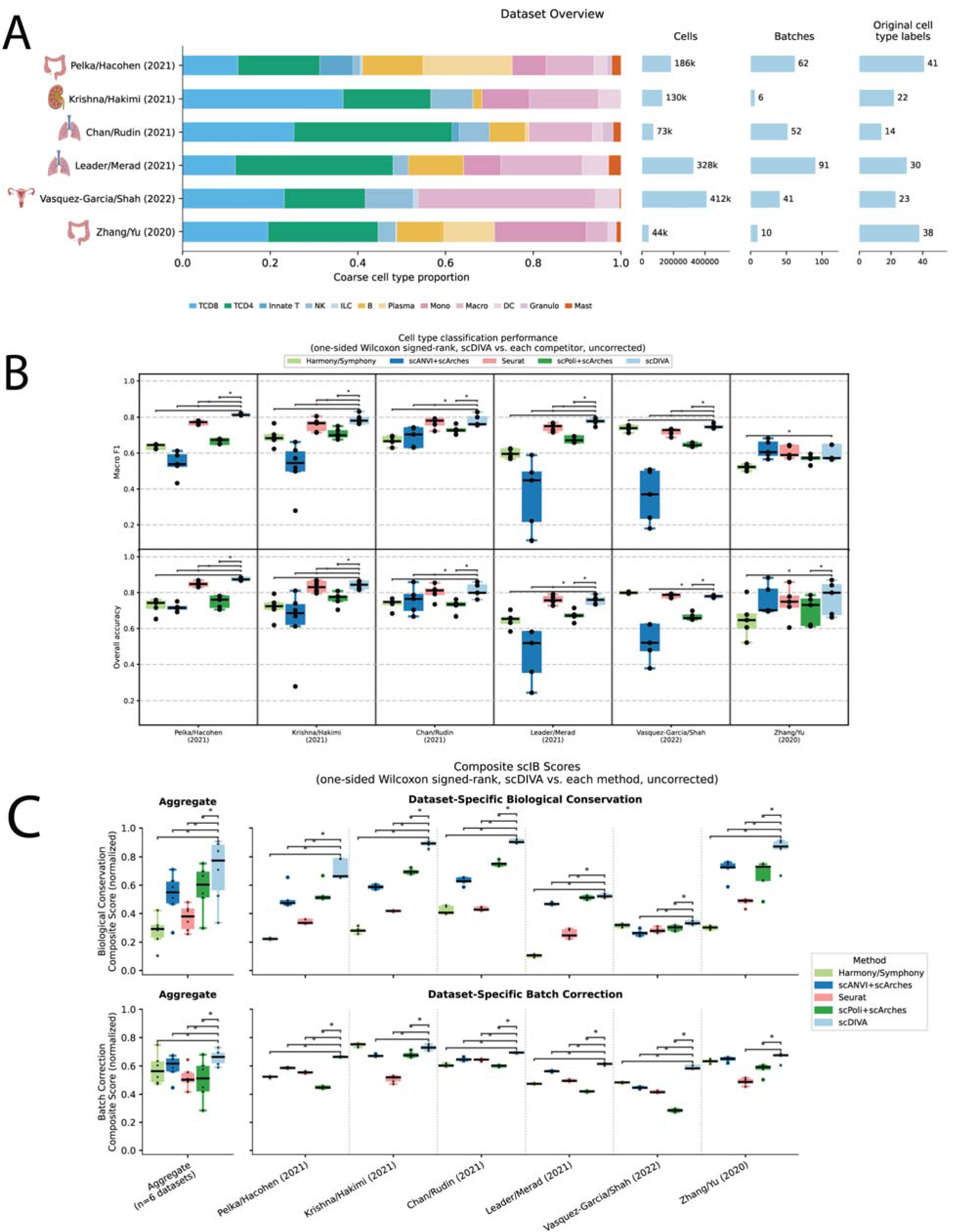
scDIVA improves fine-grained classification and integration across tumor-immune atlases. (A) Overview of the six atlases used in benchmarking. Coarse cell type composition (left), total cell count, number of batches, and number of original cell type labels (right). Atlases cover all major immune compartments (T/NK, B/plasma, and myeloid) and a wide range of cell states, spanning 14 to 41 annotated cell types per atlas, ensuring the comparisons span the diversity of the tumor immune microenvironment. (B) Macro-F1 (top) and overall classification accuracy (bottom) for scDIVA and four reference-mapping approaches: Harmony with Symphony query mapping, scANVI with scArches, Seurat label transfer, scPoli with scArches, across six pan-immune tumor atlases, evaluated on matched reference/query splits. Boxes show the distribution across splits; points are individual splits. Asterisks denote one-sided Wilcoxon signed-rank tests of scDIVA against each competitor (uncorrected, *p < 0.05). (C) Composite scIB scores for biological conservation and batch correction, aggregated globally across atlases (left) and per-atlas (right), for all five methods. Each composite is the mean of per-metric min-max-normalized scores within its category. Asterisks denote one-sided Wilcoxon signed-rank tests of scDIVA against each competitor (uncorrected, *p < 0.05).

Among these methods, scDIVA achieved the highest mean macro-F1 in five of the six atlases and significantly exceeded all four competitors in four of them (Pelka/Hacohen, Krishna/Hakimi, Leader/Merad, and Vázquez-García/Shah; one-sided Wilcoxon, p = 0.016-0.031), while outperforming Harmony/Symphony in every atlas (**Fig. 2B**, top: full win/tie/loss scorecard in **Fig. S3**). On Pelka/Hacohen, scDIVA achieved a mean macro-F1 of 0.815 against 0.769 for the next-best method, Seurat (median paired difference +0.040), and on Leader/Merad of 0.775 against 0.743 for the same competitor (+0.031 median paired difference). This advantage came largely from the cell states that competitors failed to resolve rather than the abundant populations every method recovered. For example, scANVI’s recall collapsed to 0.12 on NK cells and 0.05 on cDC2 in Leader/Merad, and Seurat’s recall to 0.08 on the Clearing.M macrophage state in Vázquez-García/Shah, while scDIVA successfully recovered each cell type (NK and Clearing.M above 0.85; cDC2 at 0.77 precision and 0.77 recall). Here macro-F1 is sensitive to these differences because it weights every class equally, while overall accuracy can obscure weak performance on smaller populations.

Vázquez-García/Shah was the only atlas where scDIVA had lower overall accuracy than a competitor (Harmony, **Fig. S3**) and tied with a second method (Seurat, **Fig. S3**) while still maintaining superior macro-F1 to both. Of the ten most abundant cell types in this atlas, scDIVA’s mean per-class F1 was marginally behind Seurat’s (−0.016, behind on eight of ten), so it was not the abundant populations that scDIVA classified more accurately. Yet across all cell types, scDIVA’s macro-F1 led Seurat by +0.034, a margin derived almost entirely from states that Seurat collapsed into a different cell type but that scDIVA resolved, such as the mid-abundance Clearing.M macrophage population (per-class F1 +0.77 over ∼12,000 cells), with smaller recoveries on M2.COL1A1, ILC, and pDC. Overall, scDIVA’s advantage was not restricted to rarer populations. Pooled across all six atlases, the per-class F1 gain over the best competing method showed no inverse dependence on a cell type’s query frequency (Spearman ρ = +0.031, p = 0.69, n = 168; **Fig. S4**); the trend was weakly positive, and within individual atlases it ranged from weakly negative to positive. Therefore, scDIVA can resolve states across a wide range of observed frequencies, spanning rare and common classes.

scDIVA again reported the highest mean macro-F1 on the Chan/Rudin atlas, though here it tied Seurat (p = 0.219). Interestingly, on macro-recall, scPoli edged out scDIVA, but this lead was due to over-calling, as scPoli had high recall but low precision on regulatory T cell predictions, for a per-class F1 of 0.45 against scDIVA’s 0.65. Since macro-F1 penalizes for the false positives that macro-recall ignores, scDIVA led scPoli significantly based on macro-F1 (0.781 vs 0.730, p = 0.031, **Fig. S3**).

Overall accuracy, as a secondary metric for classification performance, told a complementary story. scDIVA reported the highest or tied-for-highest mean in five of six atlases but significantly exceeded all four competitors in only two: Pelka/Hacohen and Krishna/Hakimi (**Fig. 2B**, bottom; **Fig. S3**). The contraction from wins in four atlases to two is notable: since abundant populations dominate the metric, the paired tests can no longer distinguish between methods, even where scDIVA’s mean performance was still higher. However, there was only one comparison in the entire overall accuracy scorecard where scDIVA lost to a competitor, against Harmony on Vázquez-García/Shah (**Fig. S3**).

scDIVA’s integration performance was its cleanest win. On the biological-conservation composite, defined uniformly across all six atlases as the mean over four metrics (cell type ASW, cLISI, ARI, and NMI^45^; see **Methods**), scDIVA significantly exceeded all four benchmarked methods in all six atlases and in the global aggregate (**Fig. 2C**, **Fig. S5**). On batch correction, scDIVA led globally against all four competitors (p = 0.016-0.031), with only a few exceptions: on Krishna/Hakimi, Harmony corrected batch more effectively, and scDIVA tied Harmony and scANVI on Zhang/Yu (**Fig. S3**). The composite lead did not come from topping every metric; on most atlases, scDIVA ranked below the best competitor on batch ASW, iLISI, and cell type ASW. The composite lead instead came from consistency across metrics, as scDIVA was highest or near-highest on the clustering metrics ARI and NMI, and free from the single-metric collapses that pulled competitors down, such as Seurat’s weakness on the graph-based batch metrics iLISI and kBET (**Fig. S5**). scDIVA’s own relative weak point was isolated-label ASW, where it ranked last on Vázquez-García/Shah; the isolated-label metrics were undefined for Krishna/Hakimi, where no cell type was isolated, so these values were excluded from the composite and reported per dataset (**Fig. S5**).

### Differential abundance testing on scDIVA embeddings identifies out-of-reference cell states

Because scDIVA isolates cell type variation in *z_y_* independently of batch, differential abundance testing on the *z_y_* embeddings reflects genuine differences in cell state composition rather than technical variation. We leveraged this disentangled embedding by coupling scDIVA with Milo^26^, treating membership in the reference or query set as the experimental condition and performing a one-sided differential abundance test for neighborhoods enriched in the query relative to the reference (**Fig. 3A**). This test makes two outcomes statistically equivalent but biologically distinct: a query-enriched neighborhood with no reference cells marks a state the reference does not contain, whereas a query-enriched neighborhood that still contains reference cells represents a shared state that is significantly more abundant in the query. The differential abundance statistic, spatialFDR^26^, flags both cases as significant, and we used the presence or absence of reference cells within the enriched neighborhoods to distinguish between them, classifying a cluster as out-of-reference when at least 25% of its query-enriched neighborhoods were reference-free, containing no reference cells (**Methods**).

**Figure 3.**
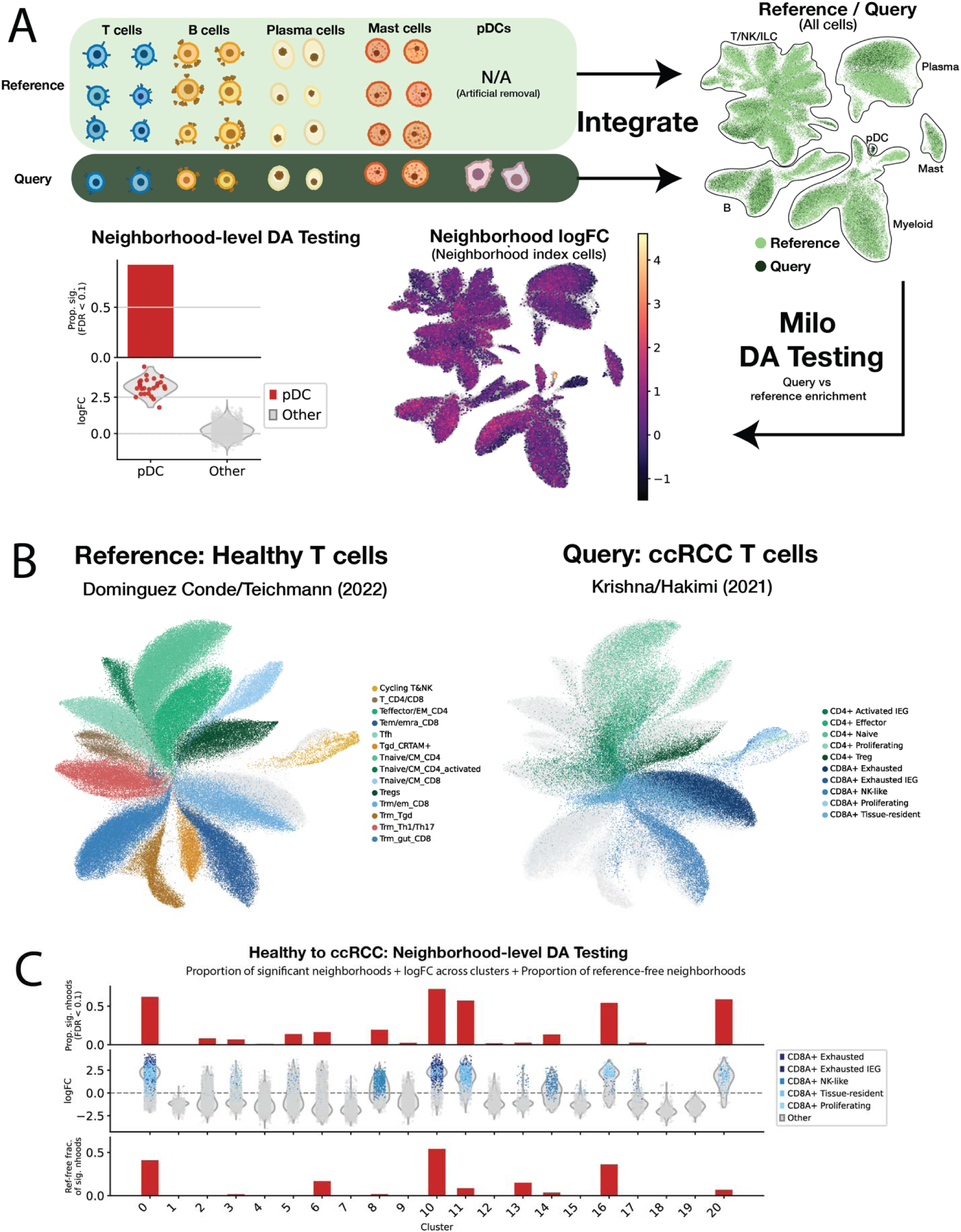
Coupling scDIVA with Milo identifies out-of-reference cell states. (A) Controlled holdout design for OOR detection. Top left: Plasmacytoid dendritic cells (pDCs) are removed from the reference of the Pelka/Hacohen CRC atlas and retained only in the query. All other cell types are shared between reference and query. This is the demonstrated example: three additional held-out populations are shown in **Fig. S7-9**. Top right: UMAP of *z_y_* embeddings from the combined reference and query containing all cells. Bottom center: Milo neighborhood log-fold change in query-versus-reference abundance, projected onto neighborhood index cells. Bottom left: Neighborhood-level differential abundance testing results for the pDC holdout compared to all other cells. Held-out pDCs are query-enriched in 80.8% of their neighborhoods, against a near-zero background in all other cells. Additional results from this held-out experiment are shown in **Fig. S6**. (B) UMAP of scDIVA cell type embeddings (*z_y_*) of the cross-context mapping of reference and query atlases. Left: Reference T cell compartment of the Dominguez Conde/Teichmann pan-tissue atlas, colored by ground truth cell type with query cells in grey. Right: Query T cell compartment of the Krishna/Hakimi ccRCC tumor-immune atlas, colored by ground truth cell type with reference cells in grey. (C) Per-cluster neighborhood-level differential abundance for the healthy-to-ccRCC mapping. Top: proportion of query-enriched neighborhoods (spatialFDR < 0.1) per Leiden cluster. Middle: neighborhood logFC per cluster. ccRCC CD8 cell states are colored, with exhausted CD8 emphasized and all other cells in grey. Every query-enriched cluster is CD8-associated. Bottom: the fraction of each cluster’s query-enriched neighborhoods that are reference-free (contain no healthy reference cell). High values mark OOR states (exhausted CD8), low values mark differentially abundant, but shared states (tissue-resident CD8).

We first tested the OOR detection sensitivity on controlled hold-outs in the Pelka/Hacohen CRC atlas by removing a known population from the reference while retaining it in the query. After artificially removing plasmacytoid dendritic cells (pDCs), a lineage transcriptionally distinct from other cell states in this atlas, DA testing flagged the held-out population as enriched in 80.8% of the neighborhoods in which these cells appeared. Extending this design to additional held-out populations bounded this method’s sensitivity (**Fig. S7-9**). We found that OOR detection sensitivity increased with transcriptional distinctness from reference states. Held-out exhausted CD8 T cells (CD8+CXCL13+) were detected as OOR nearly as strongly as pDCs, whereas GZMK+ effector-memory CD8, which are related to several CD8 effector and memory states in the reference, were flagged as OOR more weakly, with its DA signal diffused across clusters. Therefore, this framework for OOR discovery most reliably detects populations that are fully distinct from any cell state in the reference.

We then applied scDIVA across biological contexts by leveraging the T cell compartment of a healthy pan-tissue immune atlas as reference (Domínguez Conde/Teichmann^2^) and the T cell compartment of a ccRCC tumor-immune atlas (Krishna/Hakimi^39^) as the unlabeled query. The healthy reference contains no exhausted CD8 state, and naive label transfer displayed clear drawbacks: 97% of the tumor’s exhausted CD8 T cells were incorrectly assigned the nearest healthy label of tissue-resident/effector-memory CD8 (Trm/em_CD8) (**Fig. S10**). Because the healthy and tumor nomenclatures are not identical, we interpret this misassignment as the absence of a corresponding reference label rather than as a classification error, as there is no correct label in the healthy reference atlas. This misassignment, however, results from the fact that the tumor exhausted CD8 and healthy Trm/em_CD8 co-embed in *z_y_*, sharing Leiden clusters. The most enriched shared cluster was 54% tumor exhausted CD8 and 21% healthy Trm/em_CD8 (**Fig. 3B**, **Fig. S10**), consistent with the transcriptional overlap between residency and exhaustion programs in tumor CD8 T cells, which are transcriptionally similar but functionally and clonally distinct^46,47^. Other states co-embedded due to their presence in both healthy reference and tumor query datasets, such as naïve CD4 T cells and regulatory T cells (**Fig. S11-12**).

At the finer resolution of neighborhood-level DA, this framework resolves OOR cell states that are not detected purely via clustering. Five clusters were query-enriched, and the enrichment was confined to the CD8 compartment: two exhausted CD8-clusters, two tissue-resident clusters, and one proliferating cluster (**Fig. 3C**). The two tissue-resident clusters retained healthy cells in their query-enriched neighborhoods (fewer than 10% are reference-free). This suggests that the healthy tissue-resident CD8 is a normal tissue-surveillance population well-represented in the healthy pan-tissue reference and co-embedded with its tumor counterpart. However, DA testing flagged this population as differentially abundant in the tumor context. By contrast, a substantial fraction of the query-enriched neighborhoods in the two exhausted clusters were free of reference cells (41% and 54%), placing both clusters above the 25% criterion. Thus, exhausted cells represent a transcriptionally isolated population that only shares a coarse cluster with co-embedded reference cells. The proliferating cluster was also classified as out-of-reference (37% of its query-enriched neighborhoods are reference-free), as the CD8+ Proliferating population expresses high levels of activation/exhaustion markers (**Fig. S13**). The flagged cells across these five clusters express the canonical markers of terminal exhaustion (*TOX*, *PDCD1*, *HAVCR2*, *CXCL13*, *LAG3*), a program restricted to chronic antigen exposure with no healthy counterpart, whereas the tissue-resident cluster expresses residency markers (*ZNF683*, *CXCR6*) without the exhaustion signature. The genuinely tumor-restricted states were flagged as out-of-reference and the shared-but-expanded states as differentially abundant, a distinction the DA test alone cannot make.

Together, these results show that a healthy reference is insufficient to annotate immune cell states in tumors, and that scDIVA coupled with Milo flags OOR cell states that naive label transfer would otherwise misannotate. We note that OOR detection requires transcriptional distance from any reference state. Both the held-out pDCs and the tumor exhausted CD8 cells were transcriptionally isolated from all states in their respective references and therefore readily detected through DA testing.

### scDIVA integrates two independent CRC cohorts to define a conserved immune landscape

To evaluate scDIVA in a realistic cross-cohort setting, we integrated two independently generated, treatment-naïve colorectal cancer immune atlases: the fine-grained Pelka/Hacohen reference (∼186,000 immune cells, 41 immune subtypes across 62 batches) and the Zhang/Yu query (∼23,000 cells, 38 subtypes across 10 batches), restricting analysis to the tumor and adjacent-normal compartments common to both datasets. In the resulting *z_y_* embedding, cells from both cohorts co-embedded, with the major immune lineages occupying shared regions rather than separating by cohort of origin (**Fig. 4A**).

**Figure 4.**
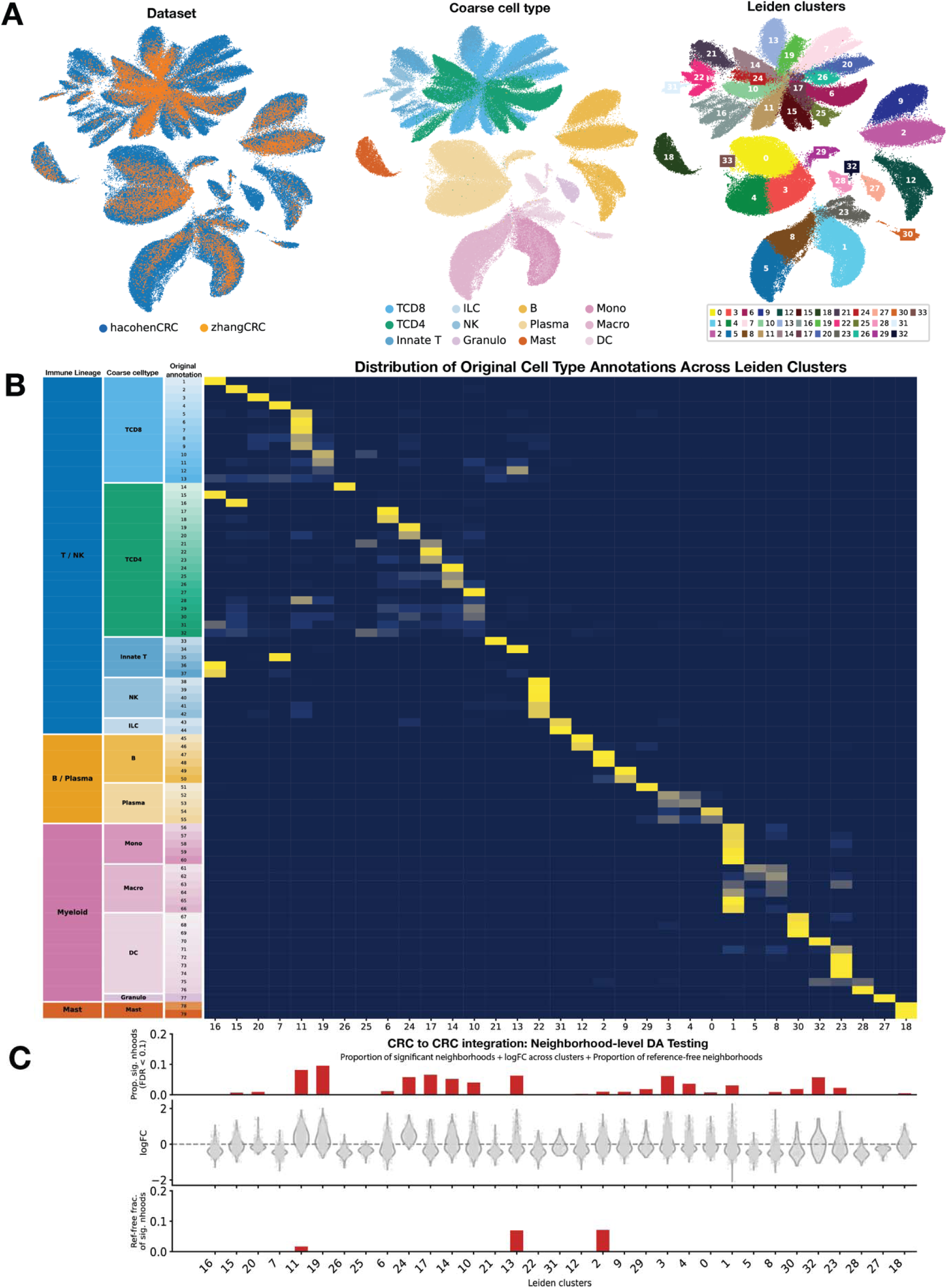
scDIVA transfers fine-grained immune annotations across independent CRC cohorts, revealing a conserved immune landscape. (B) UMAP of *z_y_* embeddings from the integrated reference (Pelka/Hacohen, ∼186,000 immune cells) and query (Zhang/Yu, ∼23,000 cells), colored by dataset of origin, coarse cell type, and leiden cluster. Peripheral blood was excluded so that both datasets share the tumor and adjacent-normal compartments. Cohorts co-embed across all lineages, indicating batch-invariant integration in *z_y_*. (C) Row-normalized abundance of the 79 cohort-specific original annotations across Leiden clusters. Fine-grained subtypes from both cohorts map onto shared clusters by lineage and coarse cell type. Because the two annotation nomenclatures are non-identical, this correspondence reflects biological coherence rather than label accuracy (within-nomenclature accuracy, Fig. 2B; full correspondence matrix, **Fig. S15**). (D) Top: per-cluster proportion of query-enriched neighborhoods (spatialFDR < 0.1) using a one-sided DA test. Middle: per-cluster distribution of neighborhood logFC distributions. Fewer than 10% of neighborhoods are query-enriched in any cluster and logFC is centered near zero, far below the held-out positive control in Fig. 3A, consistent with a conserved immune landscape. The residual enrichment localizes to cell types co-embedded across both cohorts and reflects differential abundance of shared populations rather than novel ones. Bottom: proportion of each cluster’s query-enriched neighborhoods that are reference-free to distinguish between these cases; 5 of the 502 query-enriched neighborhoods in total were reference-free.

Fine-grained cell states from the two cohorts mapped onto a shared set of Leiden clusters, with query subtypes consistently co-localizing with reference subtypes of matching lineage and state (**Fig. 4B**, **Fig. S14**). Because the two cohorts used different nomenclatures for annotation, this mapping reflected label correspondence rather than exact label accuracy (**Fig. S15,S16**). Query subtypes lacking an exact equivalent reference label distributed across similar reference states. Zhang/Yu subtypes without a dedicated reference label were annotated as the nearest available reference state: effector and effector-memory subtypes CD8-CX3CR1 and CD8-LEF1, for which the reference carries no identical label, both mapped onto the reference CD8-GZMK+ effector compartment. This is consistent with their marker expression in the blood-excluded data used here, where neither retains a naïve phenotype (**Fig. S17**). Collapsed to immune lineage, cross-cohort label transfer was essentially exact (**Fig. S16A**). The exception at coarse cell type resolution (**Fig. S16B**) was the monocyte/macrophage boundary, where 60% of Zhang/Yu macrophages were annotated as monocytes—potentially representing a difference between studies on where to define the monocyte/macrophage boundary across a continuous differentiation trajectory rather than a cell type classification error. The single cross-lineage exception was cytotoxic CD4 T cells (CD4-GNLY, a small tissue population that was mapped predominantly to the CD8+ GZMK+ effector compartment, **Fig. S15)**. Within this population, CD8A and CD8B expression were associated with the predicted label rather than the original one (**Fig. S18**), suggesting a source-annotation discrepancy from the original dataset rather than a label transfer error. These cells also express a cytotoxic effector program (*GZMK*, *PRF1,* **Fig. S17**) and overlap CD8 effector-memory cells in the embedding (**Fig. S19**).

To test whether the two CRC data sets shared a common immune landscape, we applied Milo to the integrated *z_y_* graph with reference-versus-query membership as the condition (one-sided test for query-enrichment; spatialFDR < 0.1; **Methods**). Of 20,892 neighborhoods, only 502 (2.4%) were query-enriched; no cluster exceeded ∼10% query-enriched neighborhoods (**Fig. 4C**)—roughly an order of magnitude below the held-out pDC positive control of **Fig. 3A**—and log-fold-changes were centered near zero (maximum ∼2.2), confirming that the framework reports conservation, rather than false positive OOR calls, when the underlying biology is shared.

The residual query enrichment that did occur fell on populations shared by both cohorts (**Fig. 4C**), marking differential abundance rather than novel cell states. This is clearest for GZMK+ effector-memory CD8 T cells (cluster 11; ∼8% query-enriched neighborhoods), whose reference and query populations overlap completely in the *z_y_* embedding (**Fig. S19**). Although this cell type ranked among the most query-enriched, almost all query-enriched neighborhoods retained reference cells (only 1 of 59 was reference-free). The most query-enriched cell type in this analysis, a tissue-resident CD8 population driven by CD8-CD6 and CD8-CD180, behaved identically, although its enrichment may partially reflect differences in tumor-versus-normal sampling between the cohorts rather than disease-enriched remodeling alone (**Fig. S20**).

These results from **Fig. 3** and **4** show how scDIVA distinguished between two kinds of query enrichment: (i) a population that is absent from the reference, whose query-enriched neighborhoods are reference-free, and produces large, consistent enrichment (e.g. held-out pDCs of **Fig. 3**); and (ii) a shared population that is more abundant in the query but retains reference neighbors and produces modest enrichment (e.g. cluster 11 here). The fraction of query-enriched neighborhoods that are reference-free, not the spatialFDR value alone, distinguishes between the two. Together with the hold-out experiments, these results show that scDIVA supports fine-grained immune annotation across independent clinical cohorts while flagging OOR states that are transcriptionally distinct from the reference.

## Discussion

We presented scDIVA, a semi-supervised deep variational model that performs integration, fine-grained cell type annotation, and tests for OOR cell states in a single framework. Giving batch identity its own encoder and latent subspace yields a cell type embedding that is batch-invariant by design; pairing that embedding with reference-versus-query DA testing turns the detection of OOR states into a formal, population-level test. Benchmarked against four established reference-mapping approaches across six tumor-immune atlases spanning five cancer types, scDIVA achieved the highest mean macro-F1 in five of six atlases and was also the best performer on aggregate measures of biological conservation. The DA coupling was sensitive enough to flag states with no reference counterpart, including held-out cell types in controlled experiments and tumor-specific exhausted CD8 T cell missing from a healthy reference atlas, while reporting a largely conserved landscape between independent tumor-immune atlases of the same tumor type.

The central design choice in scDIVA is the structural separation of technical and biological variation into dedicated latent subspaces, in contrast to the implicit batch conditioning of scVI, scANVI, and scPoli, or the adversarial batch removal of scDREAMER, and alongside a growing set of methods such as inVAE, biolord, DRVI, and FADVI, the last of which likewise factorizes label, batch, and residual variation into dedicated subspaces. These methods target general-purpose integration and are evaluated in settings where reference and query share cell type composition; scDIVA coupled with Milo is built for a complementary setting, as we couple the disentangled embedding with a formal test to identify query states the reference lacks. As scDIVA yields an embedding that is both optimized for cell state separation and disentangled from batch, this embedding is suited for downstream DA testing in which query enrichment reflects genuine composition differences rather than a batch effect. Benchmarking confirmed that scDIVA’s advantage is larger on macro-F1 than on overall accuracy (**Fig. 2B**) and derives from the states that competitors collapse into more abundant cell type.

Coupling the cell type embedding with DA testing directly addresses the question of whether a cell belongs to a known reference cell type or not. By contrast, per-cell approaches use softmax probabilities, prototype-based uncertainty, or cell-level rejection and can confidently misassign a novel cell to a reference type, as Dann et al. ^25^ demonstrate empirically for label-transfer uncertainty and reconstruction-error metrics—the population-level alternative, namely testing for query-versus-reference enrichment with formal error control, was established with Milo to detect OOR states against healthy atlases^25^. In scDIVA, this strategy is applied to a batch-disentangled embedding to provide accurate OOR detection.

When mapping a query ccRCC T cell immune atlas to a healthy T cell reference, scDIVA labeled 97% of tumor exhausted CD8 T cells the nearest healthy label, tissue-resident CD8, because the reference has no exhausted CD8 cell type (**Fig. S10**)—a mislabeling that DA analysis flags.

Tumor exhausted CD8 T cells share a Leiden cluster with healthy tissue-resident CD8 T cells, so clustering cannot tell them apart. At neighborhood resolution, however, many of the query-enriched neighborhoods in this cluster contain no reference cells, and the healthy cells in the same cluster fall in neighborhoods that the DA test does not flag. However, not all query-enriched neighborhoods are OOR, as a state shared with the reference but more abundant in the query will be flagged as DA. We use reference cell presence to distinguish between these cases, essentially a binarized form of Dann et al.’s neighborhood composition criterion^25^. Applied across two independent colorectal cohorts that share their immune composition, the same criterion reports few query-enriched neighborhoods and a conserved landscape, an order of magnitude below the held-out control experiment (**Fig. 4C**). Therefore, OOR detectability reflects the transcriptional distance between the query and the reference. Namely, the test flags populations unlike any cell type the reference contains and may miss a genuinely new query state that resembles a reference cell state (**Fig. S6**).

A complementary line of work approaches fine-grained immune annotation from another direction, training hierarchical classifiers on large, labeled reference atlases and applying them to query data without integration, with per-cell uncertainty flagging low-confidence assignments^48^. This sidesteps the reference-query statistical dependence introduced by integration, because the query does not alter the model. As an integration-based, transductive model, scDIVA does not escape that dependence—the query’s expression is part of model training. Rather than assuming the query lies within the reference distribution, we test for DA: flagging query states that have no reference counterpart. scDIVA’s classifier is supervised by reference labels alone, and **Fig. 4** evaluates transfer on an independently generated cohort rather than a held-out split of the same atlas. The two approaches are complementary: integration-free annotation maps cells onto an established hierarchy, while scDIVA provides reference-to-query transfer with explicit disentanglement and a population-level, FDR-controlled test for what the reference lacks.

A rapidly expanding area in the field applies single-cell foundation models pretrained on millions of cells, to produce general-purpose, transferable embeddings^28, 30^. These models have nonetheless not consistently outperformed task-specific methods on annotation and integration^35^. scDIVA, by contrast, is lightweight, trained per task, and provides two properties that these pretrained, general-purpose embeddings do not: structural separation of batch and cell type variation into dedicated subspaces, and a population-level test for OOR states.

Limitations of the current study include the dependence of scDIVA’s performance on the quality of the reference labels. The classifier cannot annotate the query more finely than the labels provided by the reference atlas^10^. An OOR state nested inside a broad reference cell type, e.g. a distinct CD8 T cell state under a generic CD8 label, may be absorbed into that coarse-grained annotation and go undetected as a novel state. We used two colorectal data sets to demonstrate integration across cohorts and confirmation that the OOR flagging does not produce widespread false positives. Additional replication across a range of atlases will help calibrate thresholds and establish generality.

Other limitations are properties of the model and workflow. As a transductive model, scDIVA is retrained from scratch to integrate or annotate each new dataset, unlike reference-mapping methods that project new data onto a fixed embedding. The single-pass design keeps each run simple, but incorporating a new query requires retraining rather than projection. The DA step is applied to the trained embedding as a post-hoc analysis rather than optimized jointly with the model, though it runs automatically as part of the scDIVA workflow. The DA test is resolved at the level of neighborhoods, each defined by a sampled subset of index cells (**Fig. 3A**), rather than as the per-cell label or confidence score that classifier-based methods produce. This neighborhood-level resolution is the cost of testing at the population level with formal error control. Finally, the reference presence criterion that separates OOR from DA states is a heuristic: the 25% threshold was chosen to separate the regimes we observed rather than calibrated against ground truth, and because it is applied per cluster it inherits the resolution of the Leiden clustering, so an OOR state confined to part of a cluster may fall below the threshold. Our DA test compares the query directly against the reference atlas without a matched control dataset; Dann et al. showed that this design can inflate false positives through residual batch effects^25^. scDIVA’s batch-invariant embedding is intended to mitigate that source of inflation, but a matched-control design and direct benchmarking against standard embeddings remain the definitive tests.

## Methods

### scDIVA model design

The scDIVA model is based on a Domain Invariant Variational Autoencoder (DIVA), a deep generative model that learns latent subspaces to disentangle variation relevant for label prediction from domain-specific variation to improve domain generalization^38^. DIVA was originally applied to image classification tasks where each labeled training example consisted of an image, a domain label, and a class label. DIVA uses a semi-supervised classification approach, so that test images with known domain but unknown class label are supplied at training time. We adapted DIVA for scRNA-seq integration and cell type classification where each labeled training example consists of a single-cell gene expression profile, a cell type label, and a batch label.

By adapting DIVA to gene expression data, we made several architectural and objective changes. The original convolutional encoders and decoder, designed for 2D image inputs, were replaced with fully connected layers suited to the flat structure of gene expression vectors. Each encoder maps the input through a hidden layer to two parallel heads that output the mean and scale of the approximate posterior, where the scale head uses a Softplus activation. The decoder reconstructs the input from the concatenated latent representations through a fully connected hidden layer followed by a ReLU output layer. Because gene expression values are continuous, unlike binary image pixels, we replaced the Bernoulli reconstruction likelihood used in the original DIVA with a continuous reconstruction penalty, the L2 norm between the input and its reconstruction summed over the minibatch. All network weights are Xavier uniform initialized with biases set to zero.

### scDIVA training process

We assume that a single sample from our data is generated by the following process:

1. Sample the label *y* and domain *d* respectively from *Cat(p_y_)* and *Cat(p_d_)*
2. Sample latent variables from continuous priors: *z_y_*∽*p*_θy_ (*z_y_*|y),*z_d_*∽*p*_θd_ (*z_d_*|d),*z_x_* ∽ p(*z_x_*)
3. Sample the observable *x* ∽*p_θx_* (*x*|z_y,_z_d,_*z_x_*)
4. Return (*y,d*,*x*)

The joint distribution factorizes as:

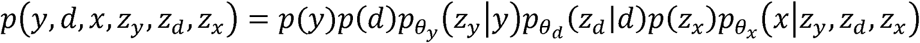

A single sample can be separated into three sources of variation: domain d can be encoded in *d*, label y can be encoded in *z_y_*, and residual variation is encoded in *z_x_*. We now approximate the posterior distribution of the latent variables with a variational posterior as:

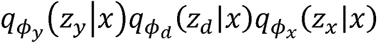

Critically, we assume label *y* and domain *d* are independently conditioned on the input example. This is reasonable for scRNA-seq data, as the batch effect (domain) information should be unrelated to cell type (label) information. By disentangling these sources of variation, examples from the same batch cluster together in *z_d_* regardless of cell type, and examples with the same cell type cluster together in *z_y_* regardless of batch. Therefore, latent spaces *z_y_* and *z_d_* encode cell type and batch information independent of other sources of variation. scDIVA then applies classifiers to these latent spaces for prediction of cell type and batch labels (**Fig. 1A**).

Successful classification of cell type and batch is dependent on the encoders, *q_фy_*(*z*_y_|*x*),,q_фd_(*z_d_*|*x*), and, q_фx_(*z_x_*|**x**) capturing information-rich embeddings of the appropriate labels. DIVA leverages a single decoder *p_θ_* (*x*|z_y,_z_d,_*z_x_*) to reconstruct the input x. To optimize the parameters for the encoders and decoder, DIVA minimzes the following variational lower bound for each input x. In this function, the first term represents the reconstruction loss and the latter terms serve as regularizers to minimize the difference between the posterior and prior distributions:

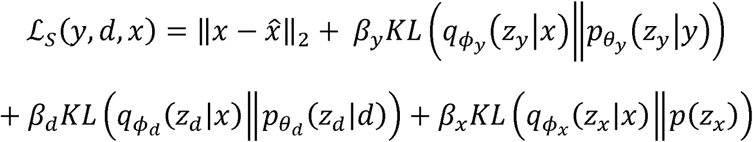

Ultimately, scDIVA uses the latent space embeddings to classify batch and cell type labels. The cross-entropy classifier loss for each of the two classifiers *q_wy_*(y,|*z_y_*) for cell type and, *q*_wd_(d.|*z_d_*) supervised training loss incorporates the variational autoencoder loss as well as the auxiliary for batch:

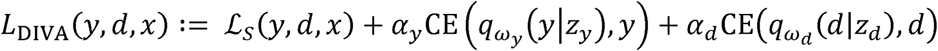

Note that DIVA is semi-supervised and utilizes batch information from the test set to further optimize batch label classification by learning the structure of the unlabeled data. In doing so, we must marginalize over the cell type label information. We enumerate all possible one-hot labels y_c_ for c = 1,…, C and weight each by the classifier’s predicted probability, *q*_wy_(y_c_; *z_y_*):

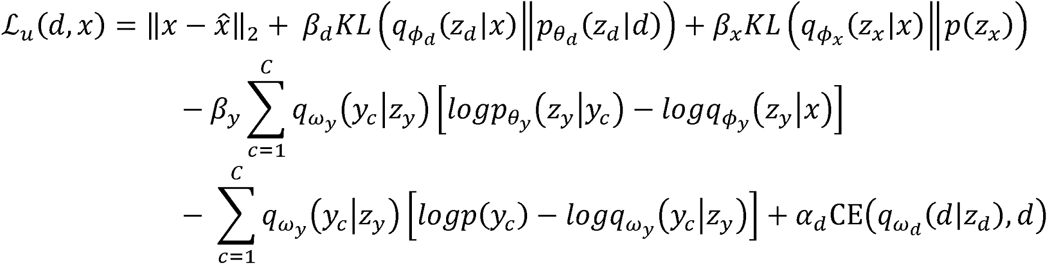

Incorporating the unsupervised loss, variational autoencoder loss, and supervised classifier loss yields a total loss function. This semi-supervised loss with N training samples (*d,x,y*) and M test samples (*d,x*) is given by:

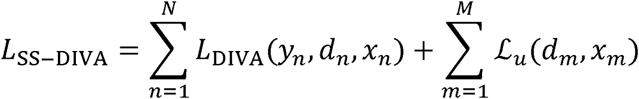

the model can experience posterior collapse. To avoid this, we warm up *β*_y_,•*β*_d_, and •*β*_z_ and over However, if the KL divergence penalty is applied at full strength from the start of the first epoch, a *t*._warmup_, which defaults to 50 epochs:

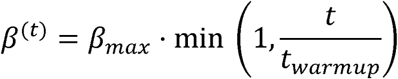

### Unsupervised batch scheduling

As a semi-supervised model, the training protocol for scDIVA balances supervised and unsupervised learning. The supervised component uses labeled data to guide training, whereas the unsupervised component uses unlabeled data to improve generalization and capture additional information about batches in the test set. We incorporate unsupervised batches periodically during training based on a predefined interval. At every training batch, we determined whether the batch should be unsupervised using the following criteria:

1. The batch index was a multiple of the predefined interval.
2. The current epoch exceeded a threshold of three epochs to ensure the model had sufficient training before introducing unsupervised data.

### Preprocessing input data

Prior to training scDIVA, we remove low quality cells and log-normalize the count matrix by normalizing to 100,000 transcripts per cell then log-transforming. To define the feature set, 3000 HVGs from the reference are merged with a set of 277 immune markers derived from immune subtype annotation signatures from tumor-immune atlases^39–44^. We found that increasing the number of HVGs did not result in an increase in classification accuracy.

### OOR cell state identification with Milo

To identify immune populations in the query that are OOR, we apply Milo to the scDIVA cell type embeddings (*z_y_*) (**Fig. 3**). We followed the standard Milo workflow^26^, building a KNN To identify immune populations in the query that are OOR, we apply Milo to the scDIVA cell graph (K = 30, d = 64) on the _y_ embeddings, defining neighborhoods with makeNhoods(prop = 0.1, K = 30, refined = FALSE), aggregating cell counts per neighborhood with countCells (samples = “batch”), and computing intra-neighborhood distances with calcNhoodDistance. We used unrefined neighborhood sampling (refined = FALSE) rather than Milo’s default refined scheme, which was necessary for tractability for large-scale tumor-immune atlases. Refined sampling repositions each sampled index cell to the median profile of its neighborhood before is noisy; because scDIVA’s *z_y_* embedding already isolates cell type structure from batch and building neighborhoods, improving neighborhood compactness when the underlying embedding residual variation, index cells sampled directly from it fall within transcriptionally coherent regions, and we did not find the refinement step necessary for stable neighborhoods. Unrefined sampling has the further advantage of drawing index cells from observed cells rather than synthetic median profiles.

To test for differential abundance between the reference and query sets, we constructed a design matrix that linked each batch to its respective set, with the reference set as the baseline level such that a positive log-fold change (logFC) indicates enrichment in the query set. Because our hypothesis is directional, we tested for query enrichment one-sided, used a modified testNhoods rather than Milo’s default two-sided test (see ‘Modifications to the Milo workflow’).

Neighborhood counts were TMM-normalized with no minimum-mean filter, and the resulting one-sided p-values were corrected with Milo’s spatialFDR procedure using k-distance weighting. For each Leiden cluster we computed the fraction of its query-enriched neighborhoods (spatialFDR < 0.1) that contained no reference cell and classified the cluster as OOR when this fraction was at least 25%.

### Modifications to the Milo workflow

We made three substantial modifications to the standard Milo workflow to enable application to large-scale tumor-immune atlases; modified code is available at https://github.com/viraj-rapolu/scDIVA/tree/main/miloRVW. This is a fork of miloR v2.2.0^26^.

First, the default calcNhoodDistance computes intra-neighborhood Euclidean distances serially, which becomes prohibitively slow at scale. We replaced this with an implementation that parallelizes the distances calculations across available CPU cores (12 by default) and calculates upper-triangle distances across all neighborhoods in a single pass.

Second, the makeNhoods function can produce neighborhoods in which the index cell has fewer than k positive distances to the other cells in its neighborhood, either because the neighborhood contains fewer than k+1 cells or because some cells share identical embedding coordinates.

Milo’s spatialFDR correction weights each neighborhood’s p-value by the inverse of the distance from its index cell to its k-th nearest neighbor, so that densely sampled regions of the embedding are penalized more heavily during multiple testing correction^26^. When fewer than k positive distances are available, this lookup returns NA and the spatialFDR calculation fails for the affected neighborhoods. We therefore clamped the k-distance to the largest available positive distance in any such neighborhood and recomputed spatialFDR across all neighborhoods using the corrected weights; the weighted Benjamini-Hochberg procedure itself is unchanged.

Third, testNhoods is two-sided by construction: it fits a negative binomial generalized linear model with edgeR::glmQLFit and evaluates the reference-versus-query coefficient with the quasi-likelihood F-test (edgeR::glmQLFTest), which has no directional form. Because our hypothesis is one-sided, as we ask only whether a neighborhood is enriched in the query relative to the reference, we implemented a directional testing path. Neighborhood counts are TMM-normalized as in the standard workflow and transformed with limma::voom, which estimates the mean-variance relationship of the log-counts-per-million and assigns each observation a precision weight; a linear model is then fit to the reference-versus-query design (limma::lmFit) and coefficient variances are shrunk toward a fitted prior by robust empirical Bayes moderation (limma::eBayes, robust=TRUE). The reported p-value is the upper-tail probability of the moderated t statistic for the query coefficient on its moderated residual degrees of freedom. These one-sided p-values replace the two-sided ones as input to the spatialFDR correction, which is otherwise applied with k-distance weighting as discussed above. All differential abundance results reported here were generated with this directional path; the reported log-fold changes and p-values therefore derive from the voom-limma linear model rather than from the edgeR negative binomial GLM that Milo uses by default.

### Method comparison

We benchmarked performance on six tumor immune scRNA-seq atlases, each containing a range of lymphoid and myeloid cell states with cell type labels specific to the study. For 4 of the 6 atlases with >10 batches, we created ref/query splits by iteratively adding individual batches to the query until it comprised between 15% and 25% of the total dataset. Only batches containing more than 500 cells were added to the query to ensure sufficient representation. This approach was inappropriate for the Krishna/Hakimi and Zhang/Yu atlases which contained fewer batches. Since the Krishna/Hakimi ccRCC atlas contained multiregional samples from six patients, we generated six splits, each corresponding to one held-out patient—all other atlases generated five splits. We created query sets for the Zhang/Yu atlas using 1-3 batches each, maintaining the same guidelines of 15-25% of the total dataset but without following the 500 cell threshold.

We compared scDIVA’s cell type classification performance to that of Seurat, Harmony + Symphony, scANVI + scArches, scPoli + scArches, using the same reference/query splits and following the preprocessing suggested for each method (**Fig. 2A**). Harmony integration and Symphony query mapping were performed using Harmonypy (v0.2.0) and symphonypy (v0.2.3), following the symphonypy reference mapping tutorial. The reference was log-normalized to 100,000 transcripts per cell, and 3,000 batch-aware HVGs were selected. The reference was then subset to these genes, scaled (max_value = 10), and PCA was computed with 30 components (zero_center=False). Harmony was run directly via harmonypy on the reference PCA embeddings with batch as the covariate, bypassing symphonypy’s harmony_integrate wrapper to avoid shape incompatibilities between symphonypy v0.2.3 and harmonypy v0.2.0. The query was log-normalized to 100,000 transcripts per cell independently and projected into the harmonized reference space via symphonypy’s map_embedding function. Cell type labels were transferred from the reference to the query using symphonypy’s KNN classifier (transfer_labels_kNN) with default parameters (n_neighbors=5) to produce a joint 30-dimensional embedding.

For Seurat (v5.4.0), the reference was log-normalized, 3000 variable features were selected via variance-stabilizing transformation, and PCA was computed with 50 components. The query was log-normalized independently. Transfer anchors were identified between the reference and query using FindTransferAnchors (normalization.method=”LogNormalize”, reference.reduction=”pca”, dims=1:50). Cell type labels were transferred to the query via TransferData (dims=1:50), and query cells were projected into the reference PCA space via IntegrateEmbeddings (dims.to.integrate=1:50) to produce a joint 50-dimensional embedding.

Using scANVI (scvi-tools v1.3.3, scArches v0.6.1), we followed the scvi-tools scArches reference mapping vignette: raw counts were initially stored in a separate layer and reference was log-normalized to 10,000 transcripts per cell for batch-aware HVG selection (3,000 genes, Seurat flavor). An scVI model was trained on the reference using the raw counts layer with batch as a covariate and default hyperparameters. An scANVI model was then initialized from the trained scVI model (unlabeled_category=“Unknown”) and trained for 20 epochs with 100 samples per label. The query was subset to the reference gene set, and raw counts were stored in a separate layer prior to log-normalization. Query cells were prepared and loaded into the reference model via SCANVI.prepare_query_anndata and SCANVI.load_query_data, then fine-tuned for 100 epochs (weight_decay=0.0, check_val_every_n_epoch=10). Cell type labels were predicted using the fine-tuned query model’s predict function. Latent representations for reference and query cells were also extracted through the fine-tuned query model to produce a joint 10-dimensional embedding.

With scPoli (scArches v0.6.1), HVGs were identified from a log-normalized copy of the reference using batch-aware selection (3000 genes, Seurat-like), and the raw count matrix was subset to these genes. The reference model was trained using a negative binomial reconstruction loss with a condition embedding dimensionality of 5 and prototype-based learning (n_epochs=50, pretraining_epochs=40, eta=5), with early stopping on validation prototype loss (patience=20, reduce_lr=True, lr_patience=13, lr_factor=0.1). Both reference and query were subset to the intersection of their gene sets. Query cells were loaded into the reference model with labeled_indices set to an empty list and trained for 50 epochs (pretraining_epochs=40, eta=10) without early stopping. Cell type labels were assigned using scPoli’s prototype-based classifier with scaled uncertainties. 10-dimensional latent representations for both reference and query cells were extracted from the fine-tuned query model using the posterior mean. For scANVI and scPoli, all cell type labels in the query set were set to “Unknown” prior to query mapping to prevent label leakage.

We evaluated classification performance with three metrics calculated per reference/query split. Macro-F1, our primary measure, is the unweighted mean across cell types of each type’s F1 score. As the mean of per-class F1, not the F1 of macro-averaged precision and recall, macro-F1 weights every cell type equally and is insensitive to class imbalance. Overall accuracy, reported as a secondary measure, is the fraction of correctly labeled query cells and is dominated by abundant populations. Macro-recall, the unweighted mean of per-class recall, is reported in the supplement as a third measure.

To compare embeddings across all five methods, we used scIB (v1.1.7) and scIB-metrics (v0.5.3)^45^. We applied scIB to the combined reference and query embeddings generated from each dataset-specific ref/query split, evaluating each method on a consistent basis. This allowed for a standardized assessment of biological conservation and batch effect correction performance across the full integrated embedding space. Since several methods, including scDIVA, produce low-dimensional embeddings rather than corrected expression matrices, we excluded expression-dependent metrics including principal component regression (PCR) batch correction, trajectory conservation, cell cycle effects, and HVG overlap. The remaining metrics assessed batch effect correction and biological conservation. Batch effect correction metrics: batch average silhouette width (batch ASW), integration local inverse Simpson’s index (iLISI), k-nearest neighbor batch-effect test (kBET), and graph connectivity. Biological conservation metrics: cell type average silhouette width (cell type ASW), cell-type local inverse Simpson’s index (cLISI), adjusted rand index (ARI), normalized mutual information (NMI), isolated label average silhouette width (isolated ASW), and isolated label F1.

To ensure a fair comparison across studies, we performed a global min-max normalization for each individual metric across all methods, datasets, and splits prior to aggregation. The plotted composite scores (**Fig. 2C**) represent the mean of these individually scaled metrics within their respective categories (biological conservation and batch effect correction). For biological conservation we defined this composite identically for every method as the mean of the four conservation metrics available for all six atlases: cell-type ASW, cLISI, ARI, and NMI. Isolated-label ASW and F1 are undefined for Krishna/Hakimi, where all 22 cell types are present in every batch, so scIB designates no isolated label and both return NaN. Since this is a property of the dataset, not any method, we exclude the two isolated-label metrics from the biological conservation composite and report them per dataset in the supplement.

We used one-sided Wilcoxon signed-rank tests to assess whether scDIVA scores exceeded those of each competing method. Because the minimum attainable one-sided p-value ranges from 0.0312 at n=5 comparisons, to 0.016 at n=6 comparisons, Benjamini-Hochberg correction would suppress nearly all significant results regardless of effect size, so we report uncorrected p-values and indicate significant differences with asterisks (* = p<0.05). Significance markers are noted above the bracket connecting scDIVA to each competing method.

## Resource availability

### Lead contact

Further information and requests for resources should be directed to and will be fulfilled by the lead contact and corresponding author, Christina S. Leslie.

### Materials availability

This study did not generate new unique reagents.

### Data and code availability

The scDIVA model and milo wrappers are available at https://github.com/viraj-rapolu/scDIVA (release v1.6.0). The optimized fork of miloR (v2.2.0) used for the differential-abundance analysis is included in the miloRVW/ subfolder of that repository. scDIVA is released under the MIT license; the bundled miloRVW/ fork and milo_postprocessing_1dir.R, which adapts code from miloR’s graphSpatialFDR, retain the GPL-3 license of the upstream miloR package. All original code has been deposited at https://doi.org/10.5281/zenodo.21829084 and is publicly available as of date of publication.

This study did not generate new sequencing data. All analyses used publicly available, published scRNA-seq atlases:

- Clear-cell renal cell carcinoma, Krishna et al.^39^ – PRJNA705464; interactive data and processed objects via the Human Cell Atlas Data Explorer (https://explore.data.humancellatlas.org/projects/12f32054-8f18-4dae-8959-bfce7e3108e7)
- Colorectal cancer, Pelka et al.^40^ – GEO GSE178341; interactive data available at the Broad Single Cell Portal, study SCP1162
- Small-cell lung cancer, Chan et al.^41^ – https://data.humantumoratlas.org/publications/hta8_2021_cancer-cell_joseph-m-chan
- Non-small-cell lung cancer, Leader et al.^42^ – GEO GSE154826 (BioProject PRJNA609924); processed data and analysis code at https://github.com/effiken/Leader_et_al
- Ovarian cancer, Vázquez-García et al.^43^ – Synapse syn25569736; dbGAP phs002857.v1.p1
- Colorectal cancer, Zhang et al.^44^ – GEO GSE146771
- Cross-tissue immune healthy atlas, Domínguez Conde et al.^2^ – ArrayExpress E-MTAB-11536 (https://tissueimmunecellatlas.org)

## Acknowledgements

This work was supported by NCI award U54CA209975 to C.S.L. V.R. and W.W. were supported by the Tri-Institutional PhD Program in Computational Biology and Medicine (NIGMS award 5T32GM132083). We thank Sneha Mitra and Tyler Park for their helpful discussions and feedback throughout the course of this project.

## Author Contributions

Conceptualization, V.R., A.K., B.L., and C.S.L.; methodology, V.R., A.K., B.L., and C.S.L.; software, V.R., A.K., B.L., and W.W.; validation, V.R.; formal analysis, V.R.; investigation, V.R.; data curation, V.R.; visualization, V.R.; writing – original draft, V.R.; writing – review & editing, V.R., W.W., and C.S.L.; supervision, C.S.L.; funding acquisition, C.S.L. A.K., B.L., and C.S.L. developed the initial implementation of the model; V.R. developed scDIVA in its current form and performed all analyses reported here.

## Competing interests

B.L. is currently an employee of Bristol Myers Squibb. His contributions to this work were made entirely as a student in the Leslie lab at Memorial Sloan Kettering Cancer Center; Bristol Myers Squibb had no role in the conception, design, execution, analysis, or reporting of this study, and no part of the work was performed at or funded by the company. V.R. completed an internship at Johnson & Johnson during the period in which this work was carried out. The internship was unrelated to this study, Johnson & Johnson had no role in the study, and V.R. holds no ongoing financial, employment, or advisory relationship with the company. C.S.L. serves on the SAB and holds IP with Episteme Prognostics, unrelated to this work. The remaining authors declare no competing interests.

## Notes

https://zenodo.org/records/21829084

https://github.com/viraj-rapolu/scDIVA

## References

1. CZI Cell Science Program et al. CZ CELLxGENE Discover: a single-cell data platform for scalable exploration, analysis and modeling of aggregated data. Nucleic Acids Res. 53, D886–D900 (2025).

2. Domínguez Conde, C., et al. Cross-tissue immune cell analysis reveals tissue-specific features in humans. Science 376, eabl5197 (2022).

3. Tyler, M. et al. The Curated Cancer Cell Atlas provides a comprehensive characterization of tumors at single-cell resolution. Nature Cancer. 6, 1088–1101 (2025).

4. Sade-Feldman, M. et al. Defining T Cell States Associated with Response to Checkpoint Immunotherapy in Melanoma. Cell 175, 998–1013.e20 (2018).

5. van der Leun, A. M., Thommen, D. S. & Schumacher, T. N. CD8+ T cell states in human cancer: insights from single-cell analysis. Nat. Rev. Cancer 20, 218–232 (2020).

6. Pai, J. A. et al. Lineage tracing reveals clonal progenitors and long-term persistence of tumor-specific T cells during immune checkpoint blockade. Cancer Cell 41, 776–790.e7 (2023).

7. Chu, Y. et al. Pan-cancer T cell atlas links a cellular stress response state to immunotherapy resistance. Nat. Med. 29, 1550–1562 (2023).

8. Huang, X. et al. Opposing functions of distinct regulatory T cell subsets in colorectal cancer. Immunity 59, 145–160.e9 (2026).

9. Regev, A. et al. The Human Cell Atlas. eLife 6, e27041 (2017).

10. Yang, J., Shyr, Y. & Liu, Q. Benchmarking single-cell tumor immune atlases and application for uncovering cell states related to immunotherapy response. Genome Biol. 27, 159 (2026).

11. Li, S., Lücken, M., Marioni, J. C., Teichmann, S. A. & He, P. Toward informed batch correction for single-cell transcriptome integration. Nat. Comput. Sci. 6, 1–11 (2026).

12. Korsunsky, I. et al. Fast, sensitive and accurate integration of single-cell data with Harmony. Nat. Methods 16, 1289–1296 (2019).

13. Kang, J. B. et al. Efficient and precise single-cell reference atlas mapping with Symphony. Nat. Commun. 12, 5890 (2021).

14. Stuart, T. et al. Comprehensive Integration of Single-Cell Data. Cell 177, 1888–1902.e21 (2019).

15. Lopez, R., Regier, J., Cole, M. B., Jordan, M. I. & Yosef, N. Deep generative modeling for single-cell transcriptomics. Nat. Methods 15, 1053–1058 (2018).

16. Xu, C. et al. Probabilistic harmonization and annotation of singleÖcell transcriptomics data with deep generative models. Mol. Syst. Biol. 17, MSB20209620 (2021).

17. Lotfollahi, M. et al. Mapping single-cell data to reference atlases by transfer learning. Nat. Biotechnol. 40, 121–130 (2022).

18. De Donno, C. et al. Population-level integration of single-cell datasets enables multi-scale analysis across samples. Nat. Methods 20, 1683–1692 (2023).

19. Michielsen, L., et al. Single-cell reference mapping to construct and extend cell-type hierarchies. NAR Genomics Bioinforma. 5, lqad070 (2023).

20. Shree, A., Pavan, M. K. & Zafar, H. scDREAMER for atlas-level integration of single-cell datasets using deep generative model paired with adversarial classifier. Nat. Commun. 14, 7781 (2023).

21. Piran, Z., Cohen, N., Hoshen, Y. & Nitzan, M. Disentanglement of single-cell data with biolord. Nat. Biotechnol. 42, 1678–1683 (2024).

22. Moinfar, A. A. & Theis, F. J. Disentangling cellular heterogeneity into interpretable biological factors through structured latent representations. 2024.11.06.622266 Preprint at 10.1101/2024.11.06.622266 (2026).

23. Aliee, H. et al. inVAE: Conditionally invariant representation learning for generating multivariate single-cell reference maps. 2024.12.06.627196 Preprint at 10.1101/2024.12.06.627196 (2024).

24. Liu, W., Qu, G., Simon, L. M., Theis, F. J. & Zhao, Z. FADVI: disentangled representation learning for robust integration of single-cell and spatial omics data. 2025.11.03.683998 Preprint at 10.1101/2025.11.03.683998 (2025).

25. Dann, E. et al. Precise identification of cell states altered in disease using healthy single-cell references. Nat. Genet. 55, 1998–2008 (2023).

26. Dann, E., Henderson, N. C., Teichmann, S. A., Morgan, M. D. & Marioni, J. C. Differential abundance testing on single-cell data using k-nearest neighbor graphs. Nat. Biotechnol. 40, 245–253 (2022).

27. Weinberger, E., Lin, C. & Lee, S.-I. Isolating salient variations of interest in single-cell data with contrastiveVI. Nat. Methods 20, 1336–1345 (2023).

28. Yang, F. et al. scBERT as a large-scale pretrained deep language model for cell type annotation of single-cell RNA-seq data. Nat. Mach. Intell. 4, 852–866 (2022).

29. Transfer learning enables predictions in network biology | Nature. https://www.nature.com/articles/s41586-023-06139-9.

30. scGPT: toward building a foundation model for single-cell multi-omics using generative AI | Nature Methods. https://www.nature.com/articles/s41592-024-02201-0.

31. Hao, M. et al. Large-scale foundation model on single-cell transcriptomics. Nat. Methods 21, 1481–1491 (2024).

32. Rosen, Y. et al. Universal cell embedding provides a foundation model for cell biology. Nature 1–9 (2026) doi:10.1038/s41586-026-10689-z.

33. Mitra, S., Li, J. & Leslie, C. S. New frontiers for AI in single-cell genomics. (2026).

34. Heimberg, G. et al. A cell atlas foundation model for scalable search of similar human cells. Nature 638, 1085–1094 (2025).

35. DenAdel, A. et al. Evaluating the role of pretraining dataset size and diversity on single-cell foundation model performance. Nature Methods 23. 1447–1457 (2026).

36. Zhou, X. et al. Benchmarking zero-shot single-cell foundation model embeddings for cellular dynamics reconstruction. 2026.03.10.710748 Preprint at 10.64898/2026.03.10.710748 (2026).

37. Han, S. et al. Benchmarking single-cell foundation models for real-world RNA-seq data integration. 2026.04.17.719314 Preprint at 10.64898/2026.04.17.719314 (2026).

38. Ilse, M., Tomczak, J. M., Louizos, C. & Welling, M. DIVA: Domain Invariant Variational Autoencoders. in Proceedings of the Third Conference on Medical Imaging with Deep Learning 322–348 (PMLR, 2020).

39. Krishna, C. et al. Single-cell sequencing links multiregional immune landscapes and tissue-resident T cells in ccRCC to tumor topology and therapy efficacy. Cancer Cell 39, 662–677.e6 (2021).

40. Pelka, K. et al. Spatially organized multicellular immune hubs in human colorectal cancer. Cell 184, 4734–4752.e20 (2021).

41. Chan, J. M. et al. Signatures of plasticity, metastasis, and immunosuppression in an atlas of human small cell lung cancer. Cancer Cell 39, 1479–1496.e18 (2021).

42. Leader, A. M. et al. Single-cell analysis of human non-small cell lung cancer lesions refines tumor classification and patient stratification. Cancer Cell 39, 1594–1609.e12 (2021).

43. Vázquez-García, I. et al. Ovarian cancer mutational processes drive site-specific immune evasion. Nature 612, 778–786 (2022).

44. Zhang, L. et al. Single-Cell Analyses Inform Mechanisms of Myeloid-Targeted Therapies in Colon Cancer. Cell 181, 442–459.e29 (2020).

45. Luecken, M. D. et al. Benchmarking atlas-level data integration in single-cell genomics. Nat. Methods 19, 41–50 (2022).

46. Burn, T. N. et al. Antigen reactivity defines tissue-resident memory and exhausted T cells in tumors. Nat. Immunol. 27, 98–109 (2026).

47. Park, S. L. et al. Tissue-resident exhausted and memory CD8+ T cells have distinct ontogeny, function and roles in disease. Nat. Immunol. 27, 110–125 (2026).

48. Beltz, C. et al. Hierarchical classification of immune cell transcriptomes at population-scale. 2026.05.30.728980 Preprint at 10.64898/2026.05.30.728980 (2026).

