## Supplemental Figures for "scDIVA: semi-supervised integration and fine-grained annotation of tumor-immune single-cell atlases"

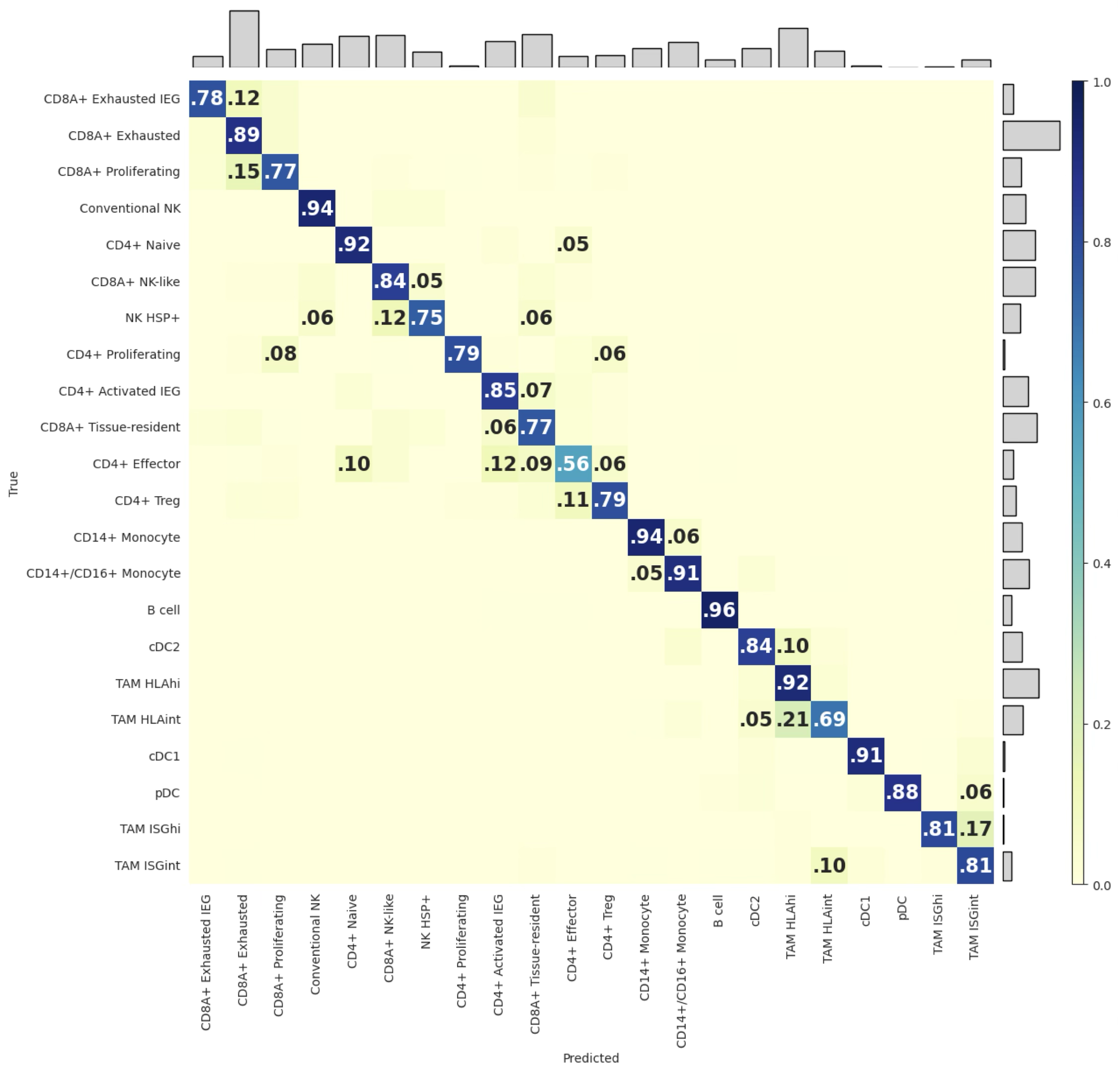

**Figure S1.** Krishna/Hakimi scDIVA confusion matrix. Related to **Fig. 1**.

True-versus-predicted cell type annotations aggregated across 6 reference/query splits.

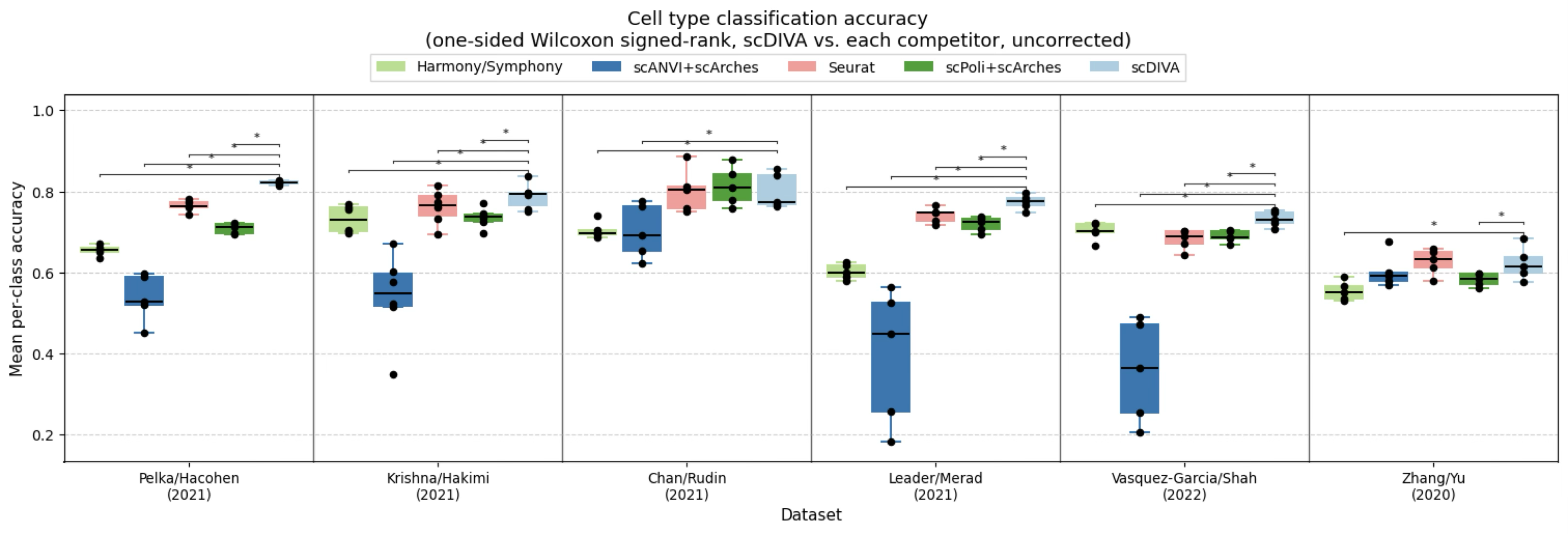

**Figure S2.** Macro recall benchmarking. Related to **Fig. 2**.

Macro recall for scDIVA and four reference-mapping approaches: Harmony with Symphony query mapping, scANVI with scArches, Seurat label transfer, scPoli with scArches, across six pan-immune tumor atlases, evaluated on matched reference/query splits. Boxes show the distribution across splits; points are individual splits. Asterisks denote one-sided paired Wilcoxon signed-rank tests of scDIVA against each competitor (uncorrected, *p < 0.05)

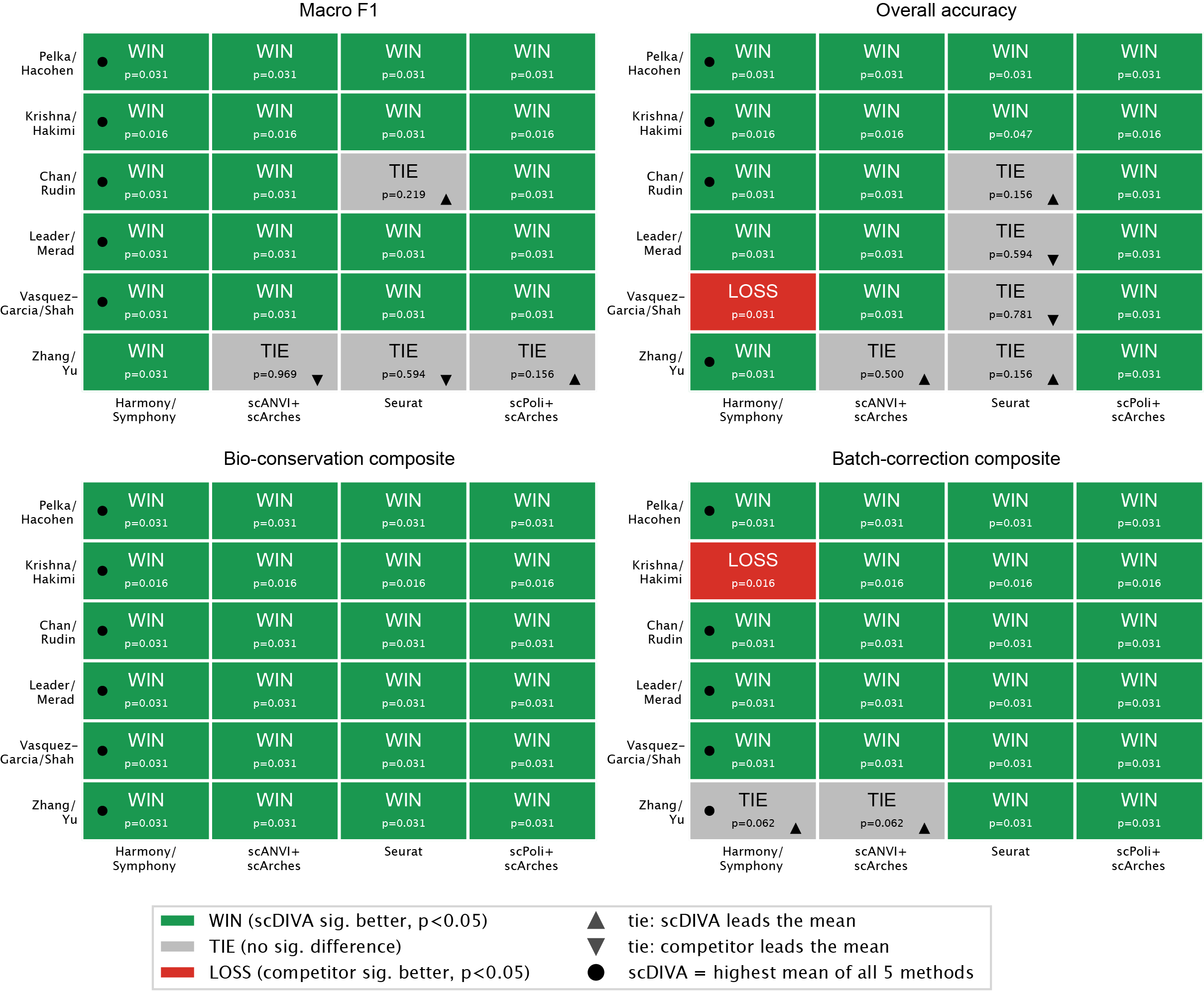
 **Figure S3.** scDIVA scorecard. Related to **Fig. 2**.

Method comparison p-values were computed using a one-sided Wilcoxon signed-rank test (uncorrected). We performed pairwise comparisons across all competitors and benchmarking atlases on macro F1, overall accuracy, biological conservation composite, and batch correction composite values. If a cell shows WIN/TIE, the p-value shown is p(scDIVA > competitor), otherwise p(competitor>scDIVA) if a cell shows LOSS. Biological composite uses the same uniform 4-metric score (cell type ASW, graph cLISI, ARI, NMI) applied identically to all atlases.

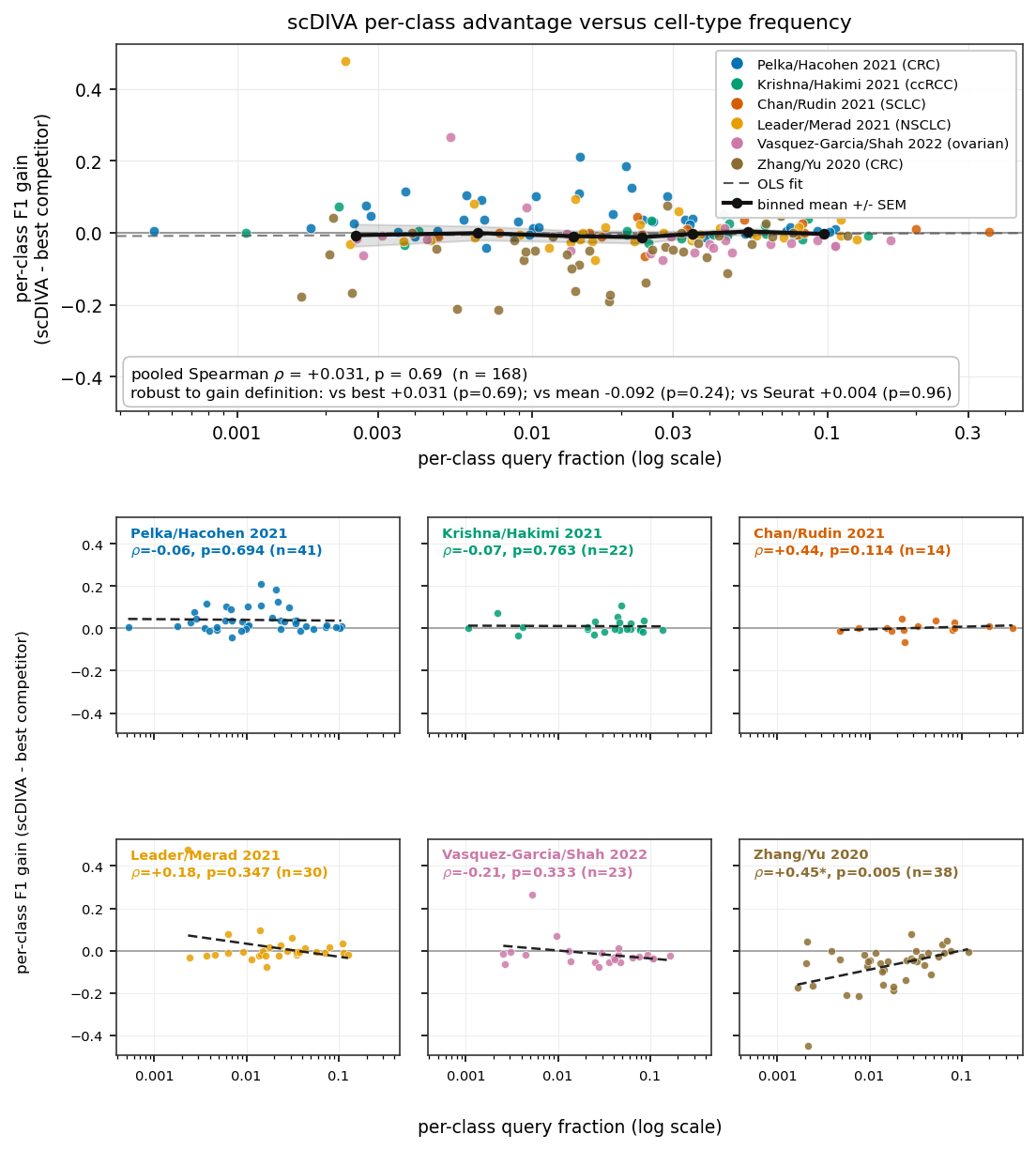

**Figure S4.** scDIVA’s per-class F1 advantage versus cell-type frequency. Related to **Fig 2.**

Per-class F1 gain (scDIVA minus best competing method) against each cell type’s query frequency, pooled across all six atlases (top; Spearman ρ = +0.031, p=0.69, n=168 cell-type-atlas pairs) and per atlas (bottom). The gain shows no negative dependence on frequency; the only significant per-class correlation is in Zhang/Yu (ρ = +0.45, p = 0.005), reflecting gains concentrated at higher frequencies in that atlas.

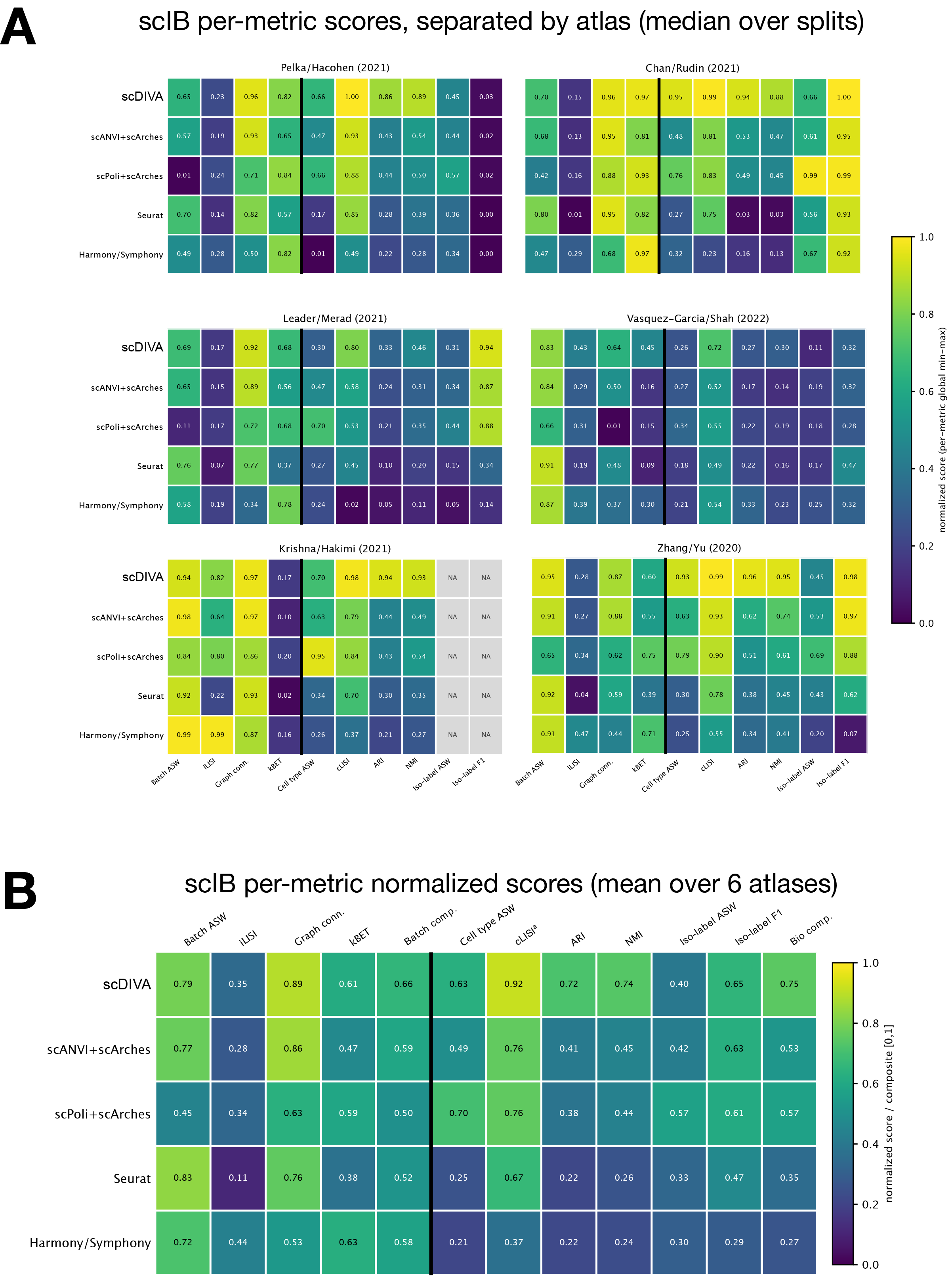

**Figure S5.** scIB per-metric normalized scores. Related to **Fig. 2**.

A) Metrics are separated by each dataset, batch effect metrics left of vertical bar, biological conservation metrics right of the bar. Since Krishna/Hakimi have all cell types present in each batch, it is not possible to compute isolated label metrics. We therefore exclude Iso-label ASW and Iso-label F1 from composite score calculation.
B) Mean of each metric across 6 atlases, including composite scores.

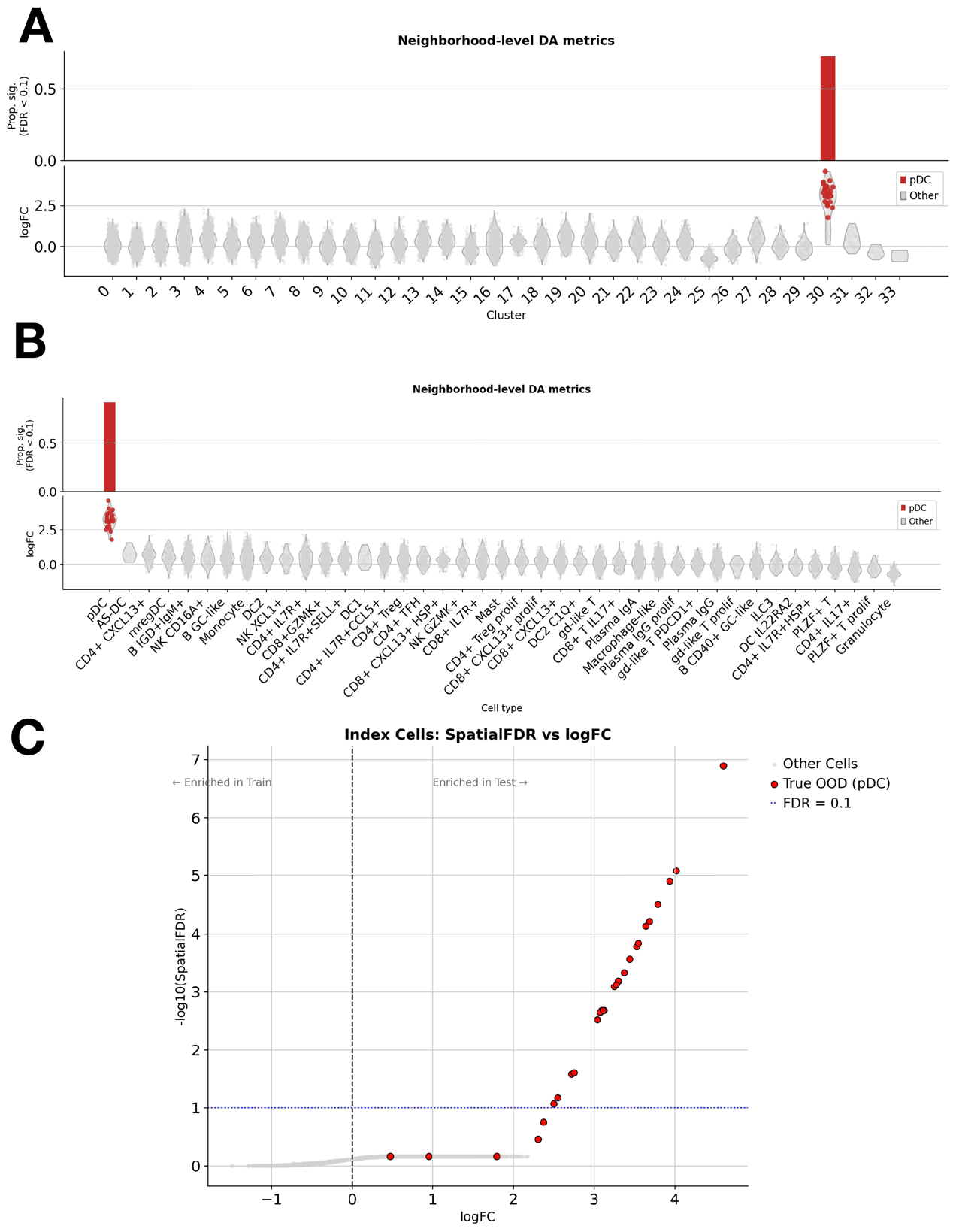

**Figure S6.** Controlled pDC holdout experiment for DA testing. Related to **Fig. 3**.

A) DA metrics grouped by Leiden cluster. Top: proportion of each group’s neighborhoods that are significantly query enriched (spatialFDR < 0.1). Bottom: neighborhood logFC.
B) DA metrics grouped by cell type. Top: proportion of each group’s neighborhoods that are significantly query enriched (spatialFDR < 0.1). Bottom: neighborhood logFC.
C) Volcano plot of -log(SpatialFDR) and logFC.

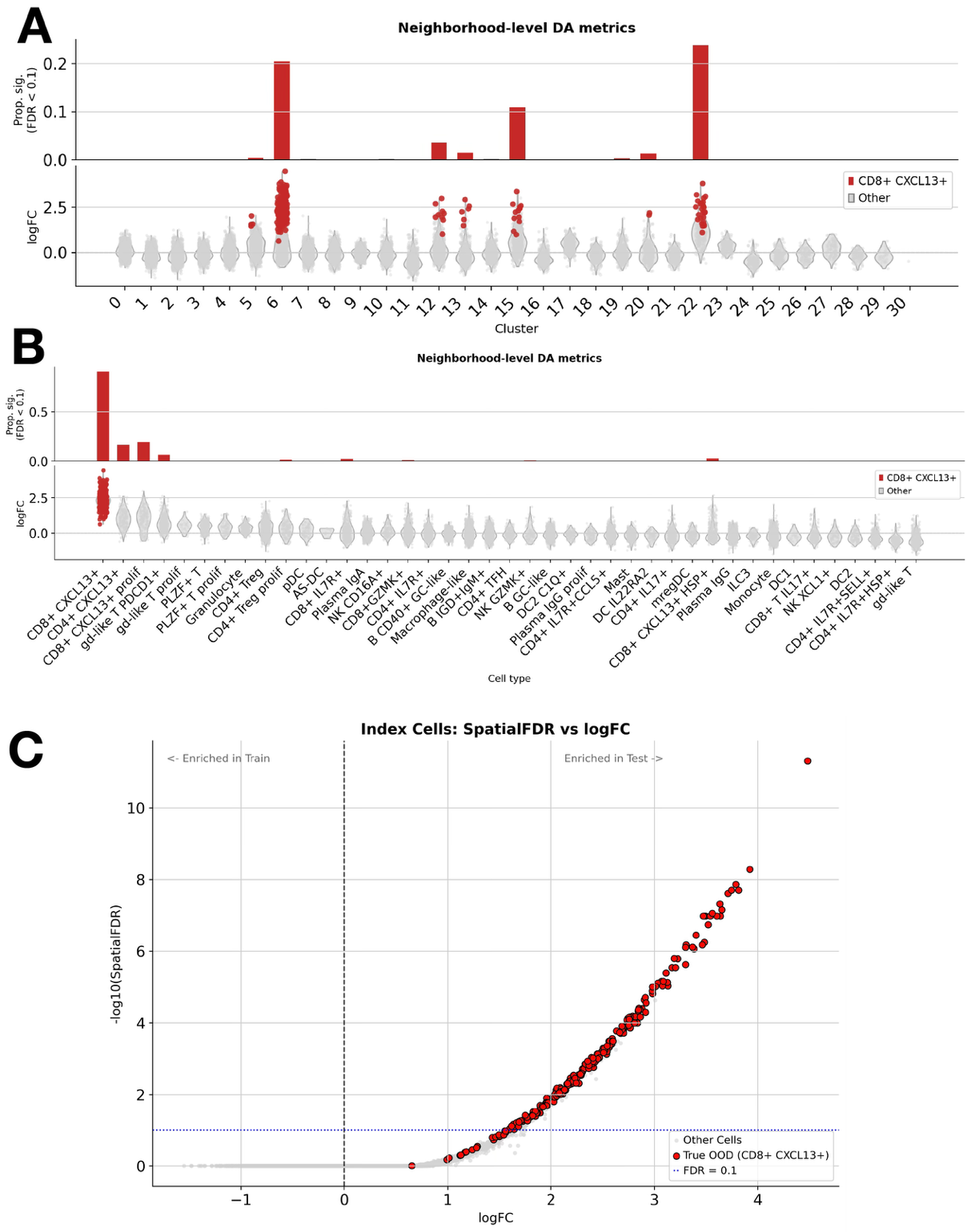

**Figure S7.** Controlled CD8+ CXCL13+ holdout experiment for DA testing. Related to **Fig. 3**.

A) DA metrics grouped by Leiden cluster. Top: proportion of each group’s neighborhoods that are significantly query enriched (spatialFDR < 0.1). Bottom: neighborhood logFC.
B) DA metrics grouped by cell type. Top: proportion of each group’s neighborhoods that are significantly query enriched (spatialFDR < 0.1). Bottom: neighborhood logFC.
C) Volcano plot of -log(SpatialFDR) and logFC.

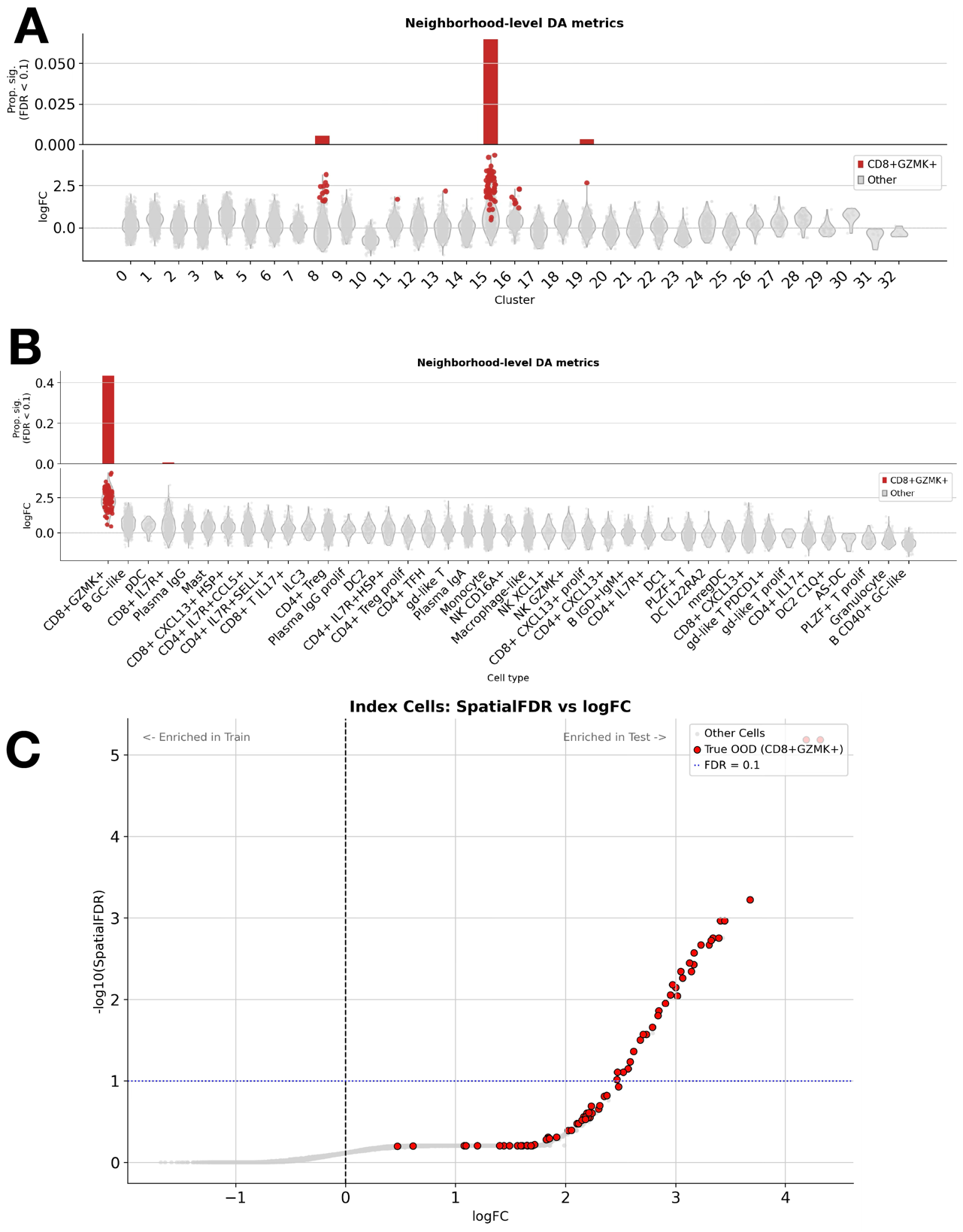

**Figure S8.** Controlled CD8+ GZMK+ holdout experiment for DA testing. Related to **Fig. 3**.

A) DA metrics grouped by Leiden cluster. Top: proportion of each group’s neighborhoods that are significantly query enriched (spatialFDR < 0.1). Bottom: neighborhood logFC.
B) DA metrics grouped by cell type. Top: proportion of each group’s neighborhoods that are significantly query enriched (spatialFDR < 0.1). Bottom: neighborhood logFC.
C) Volcano plot of -log(SpatialFDR) and logFC.

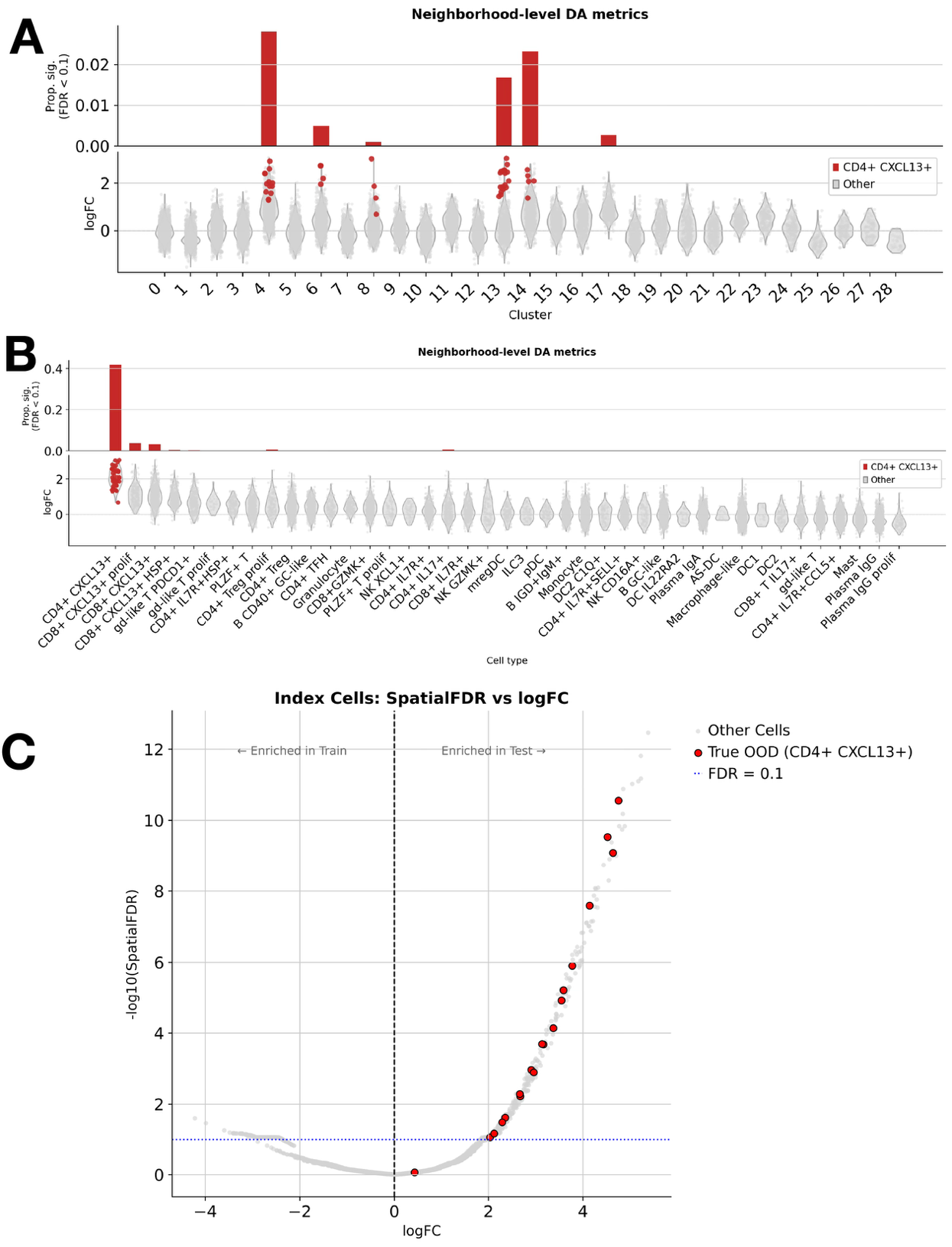

**Figure S9.** Controlled CD4+ CXCL13+ holdout experiment for DA testing. Related to **Fig. 3**.

A) DA metrics grouped by leiden cluster. Top: proportion of each group’s neighborhoods that are significantly query enriched (spatialFDR < 0.1). Bottom: neighborhood logFC.
B) DA metrics grouped by cell type. Top: proportion of each group’s neighborhoods that are significantly query enriched (spatialFDR < 0.1). Bottom: neighborhood logFC.
C) Volcano plot of -log(SpatialFDR) and logFC.

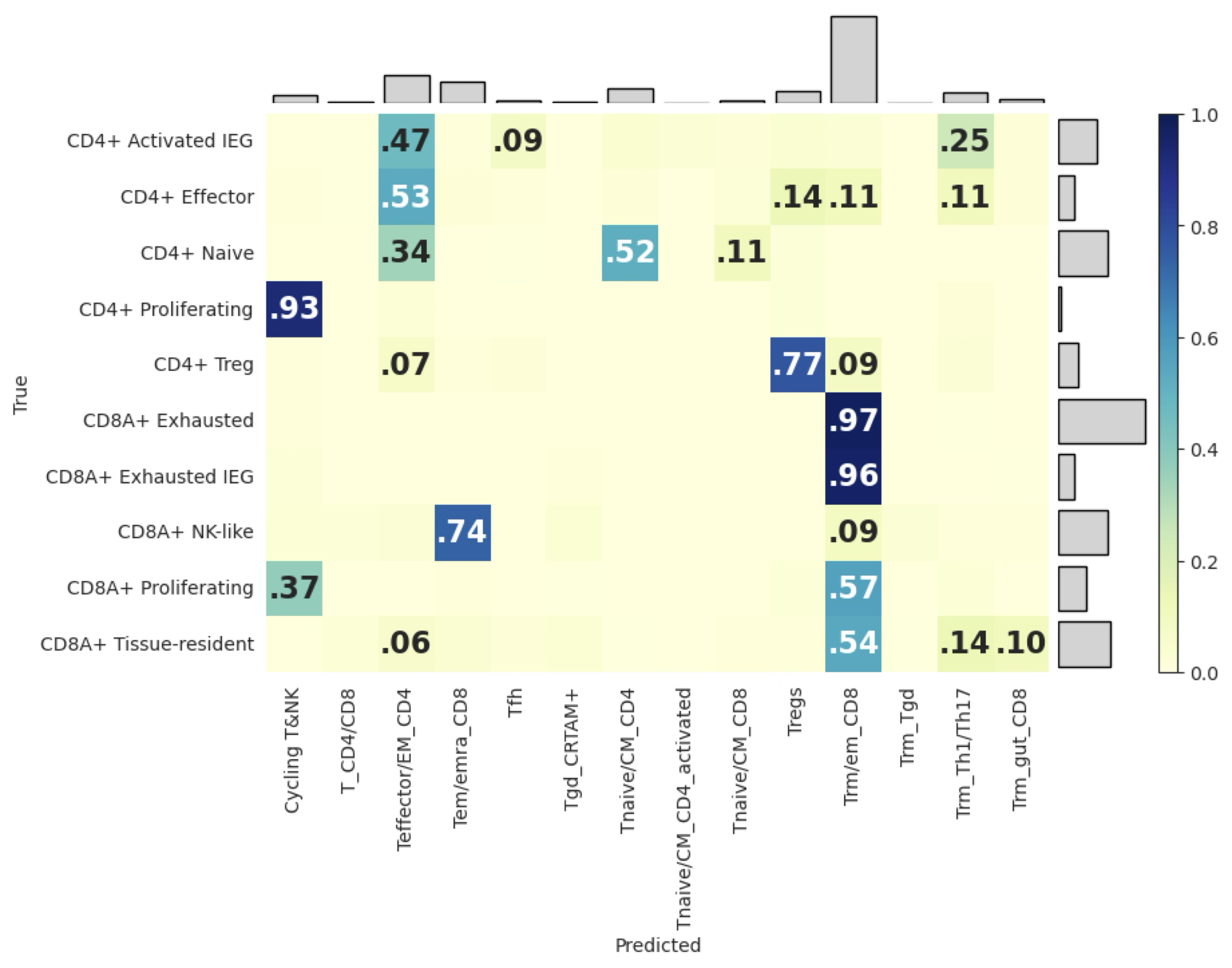

**Figure S10.** Confusion matrix from healthy to ccRCC integration. Related to **Fig. 3**.

Assignment of healthy reference labels (columns) to ccRCC query cells (rows). Because the healthy reference contains no exhausted-CD8 category, scDIVA assigns 97% of tumor exhausted CD8 T cells (and 96% of exhausted-IEG CD8) the closest available cell state, tissue resident/effector-memory CD8 (Trm/em_CD8). Since the two nomenclatures are not identical, there is no ground-truth diagonal. The confusion matrix therefore reflects the absence of a corresponding reference label rather than classification accuracy.

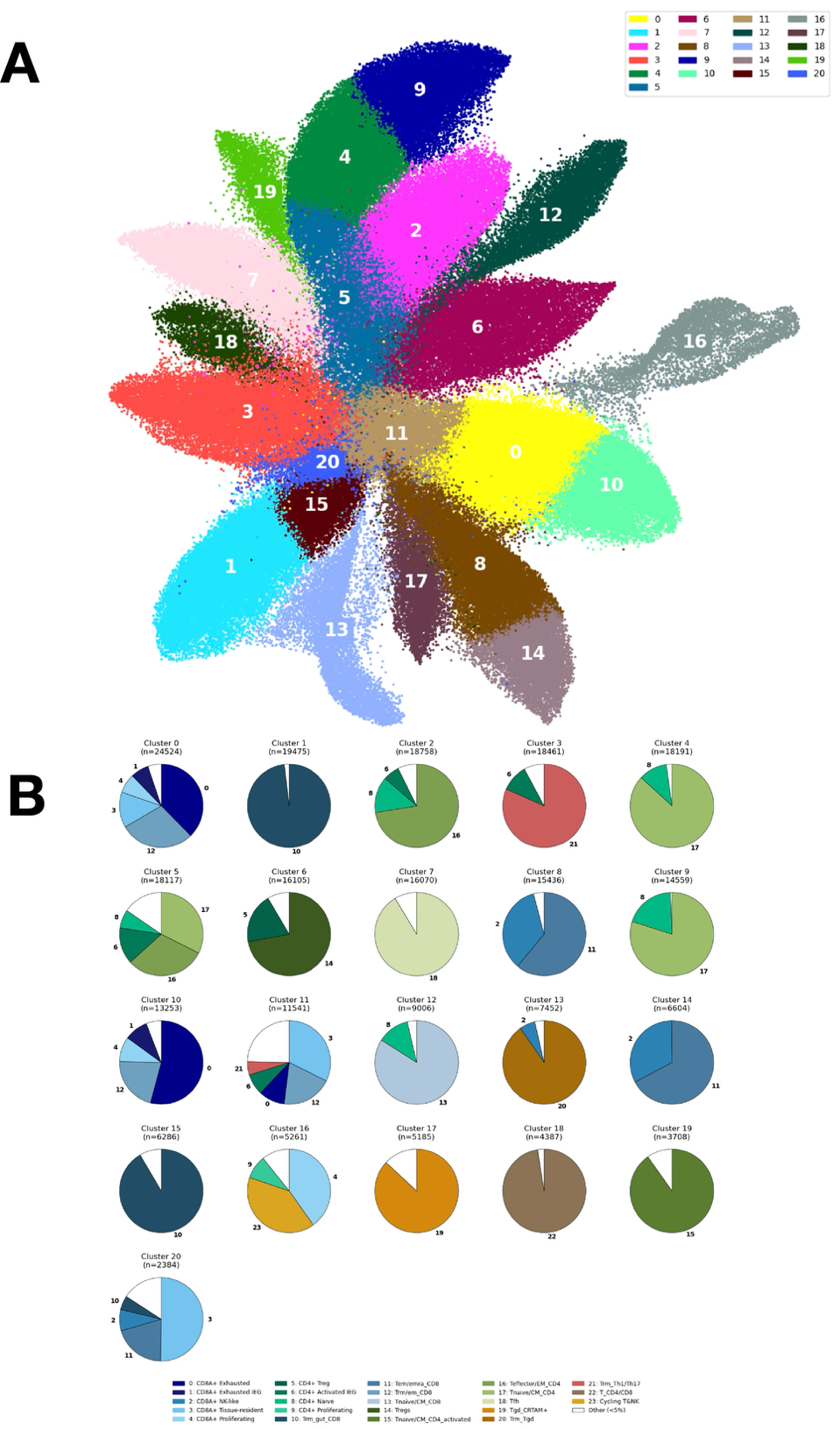

**Figure S11.** Cluster composition in healthy to ccRCC T cell integration. Related to **Fig. 3**.

A) UMAP constructed on the scDIVA zy embedding, colored by Leiden cluster.

Cluster composition. Each slice of the pie chart has a number, each corresponding to a cell type label. Any cell type that encompassed <5% of cluster composition was left blank.

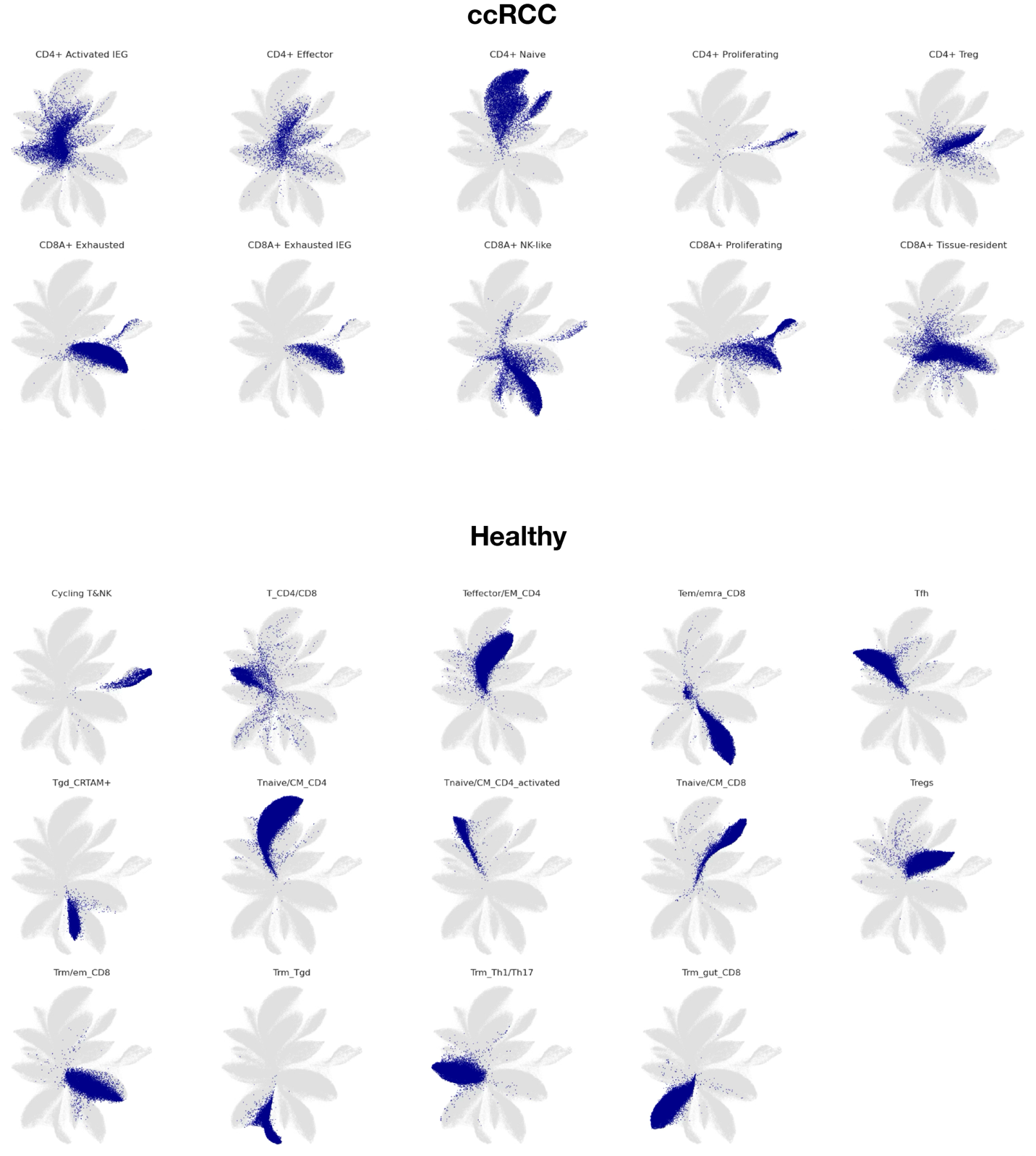
 **Figure S12.** Cell type embeddings from healthy to ccRCC integration. Related to **Fig. 3**.

UMAP of the scDIVA 𝑧_𝑦_ embedding for the combined healthy pan-tissue reference and ccRCC query T cell compartments, shown per cell type. Reference and query cells are colored separately with all remaining cells in grey. States present in both datasets, including naïve CD4 T cells and regulatory T cells, occupy overlapping regions of the embedding.

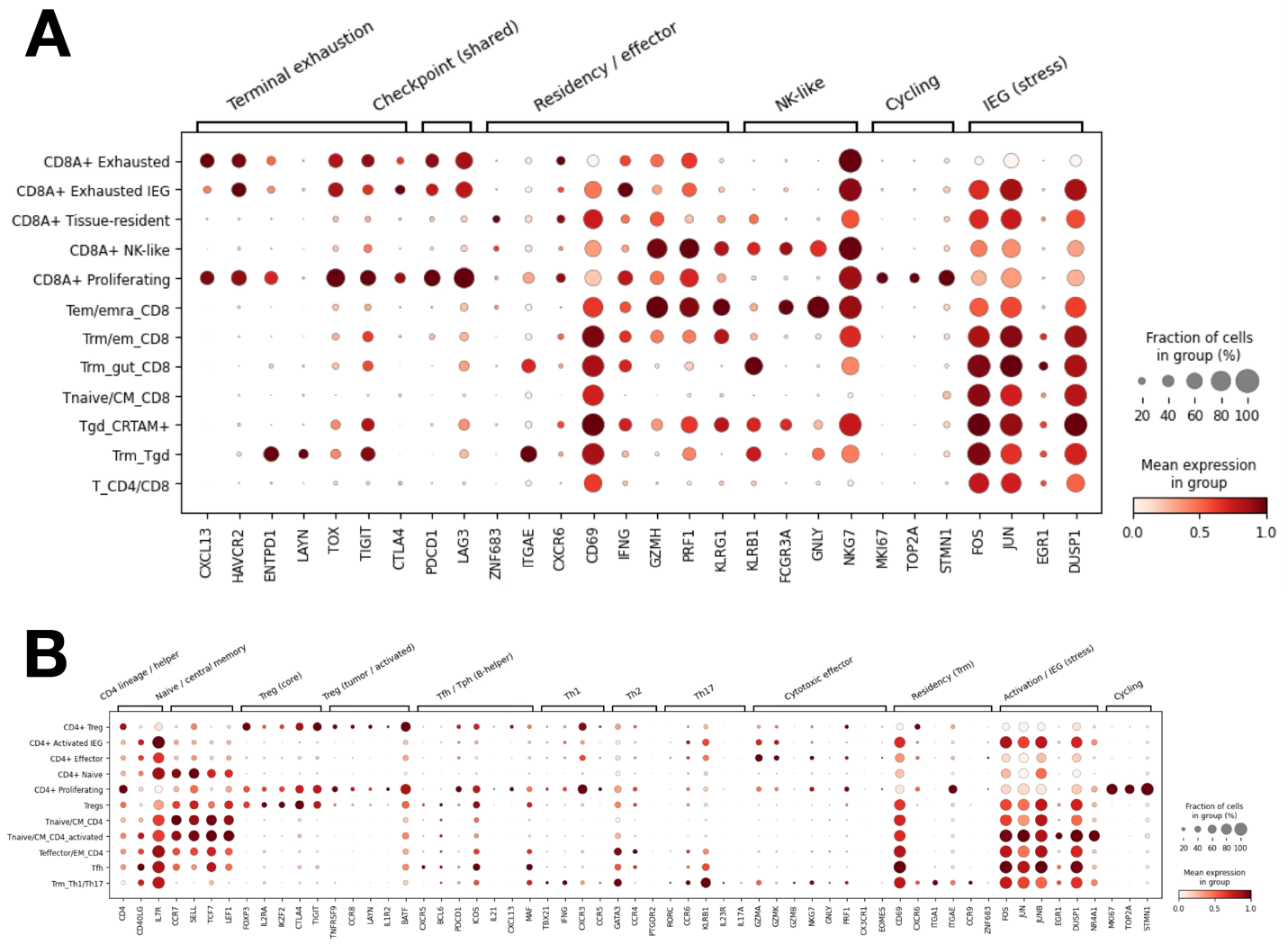

**Figure S13.** Marker gene expression in healthy to ccRCC T cell integration. Related to **Fig. 3**.

A) Marker gene expression in the CD8+ T compartment.
B) Marker gene expression in the CD4+ T compartment.

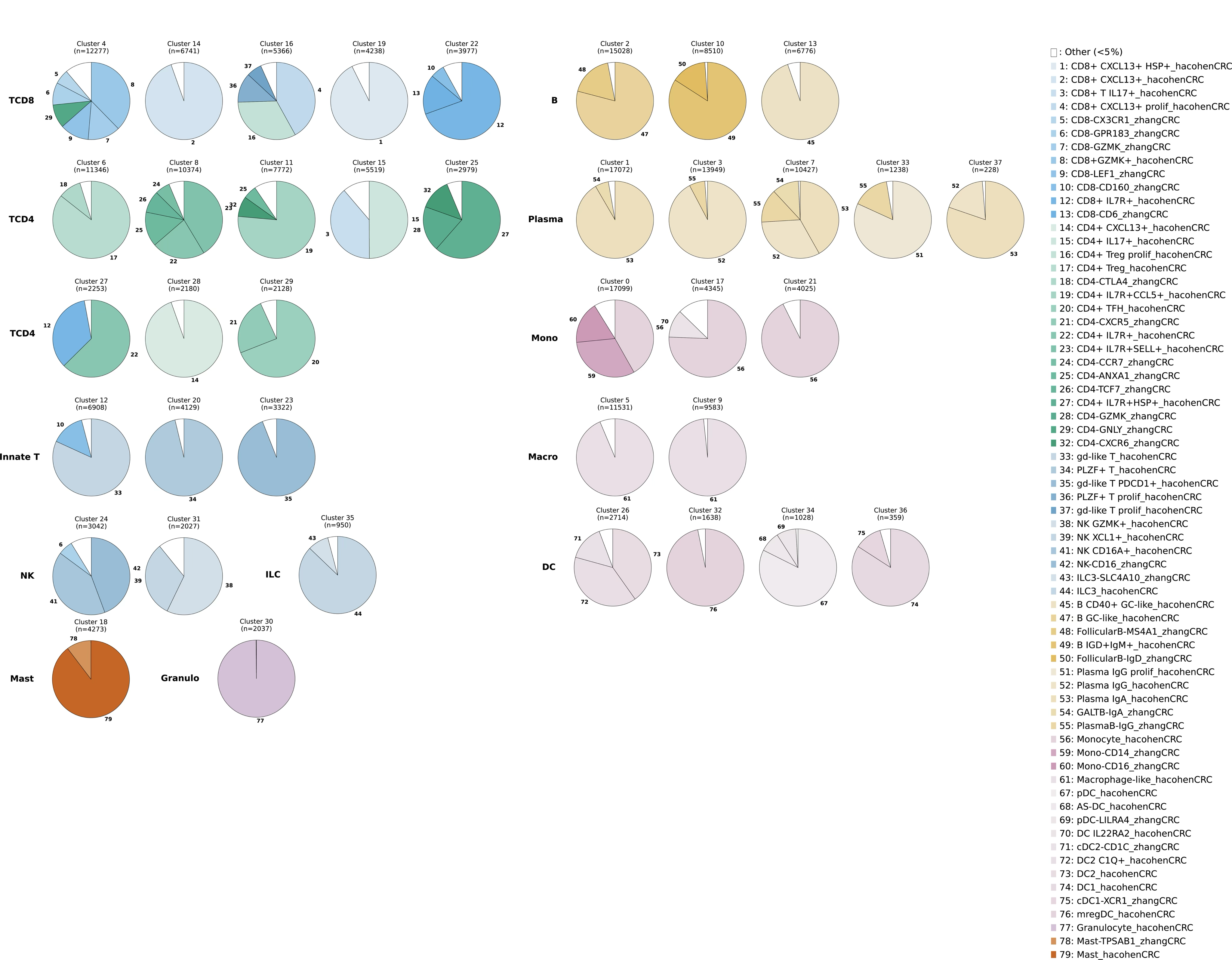

**Figure S14.** Cluster composition in CRC/CRC integration. Related to **Fig. 4**.

Each slice of the pie chart has a number, each corresponding to a cell type label. Any cell type that encompassed <5% of cluster composition was left blank. Colors correspond to the heatmap in **Fig. 4B**.

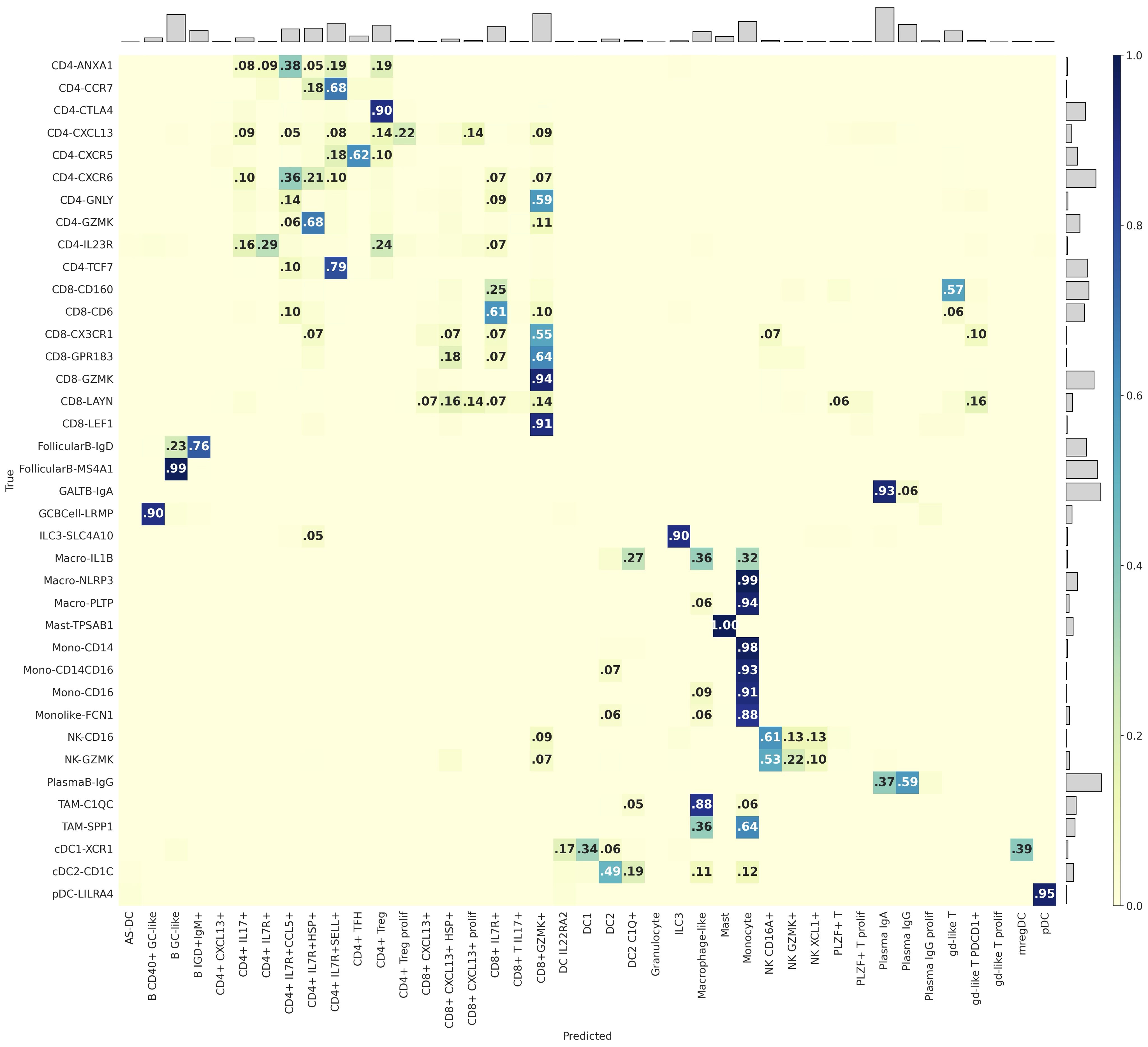

**Figure S15.** Confusion matrix from CRC to CRC integration. Related to **Fig. 4**.

Assignment of Pelka/Hacohen reference labels (columns) to Zhang/Yu query cells (rows). Most labels show good correspondence. The main exception is the myeloid compartment: the reference contains only Monocyte and Macrophage-like myeloid categories, so the finer Zhang/Yu macrophage subtypes are distributed across these two subtypes, with a majority assigned Monocyte (**Fig. S16B**).

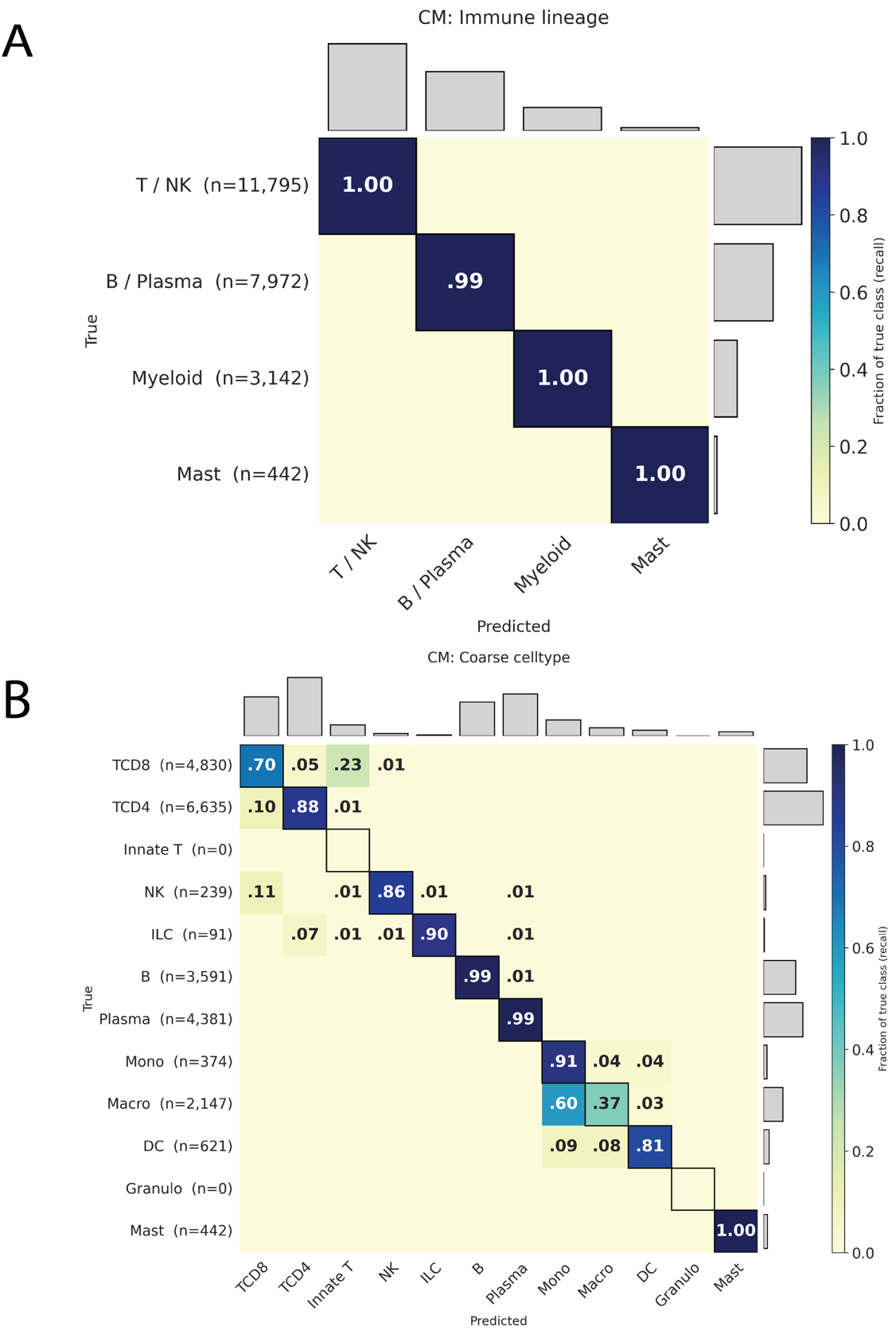

**Figure S16.** Coarse confusion matrices from CRC to CRC integration. Related to **Fig. 4**.

A) Reference annotations and query predictions from **Fig. S15** collapsed to immune lineage.
B) Reference annotations and query predictions from **Fig. S15** collapsed to coarse cell type.

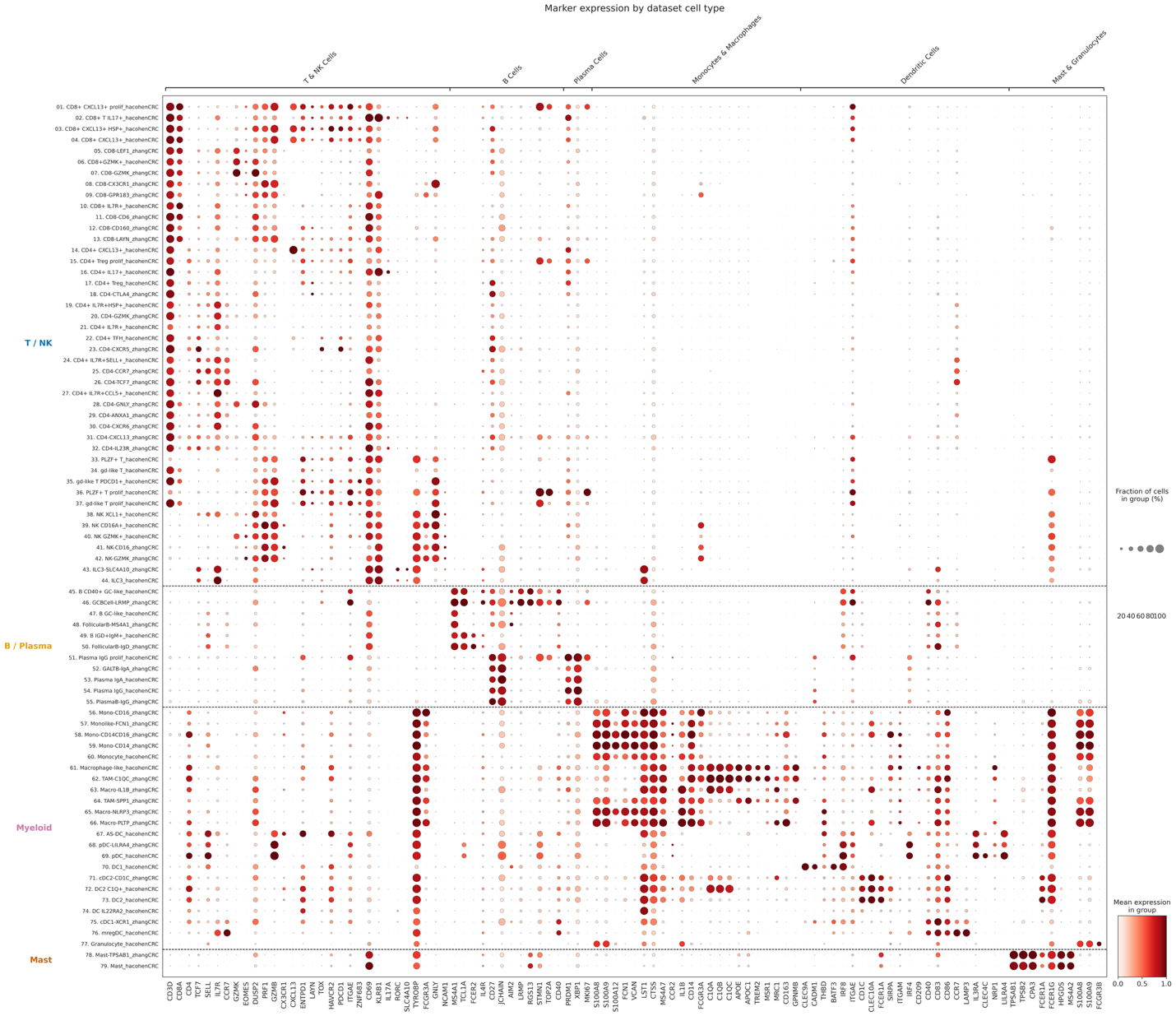
 **Figure S17.** Marker gene expression across independent CRC cohorts. Related to **Fig 4.**

Dot plot of canonical immune markers for each original annotation from the Pelka/Hacohen reference and Zhang/Yu query, grouped by immune lineage. Consistent with their assignment to the reference CD8-GZMK+ effector compartment, CD8-LEF1 and CD8-CX3CR1 express effector and effector-memory markers rather than a naïve program in the blood-excluded data used in this integration.

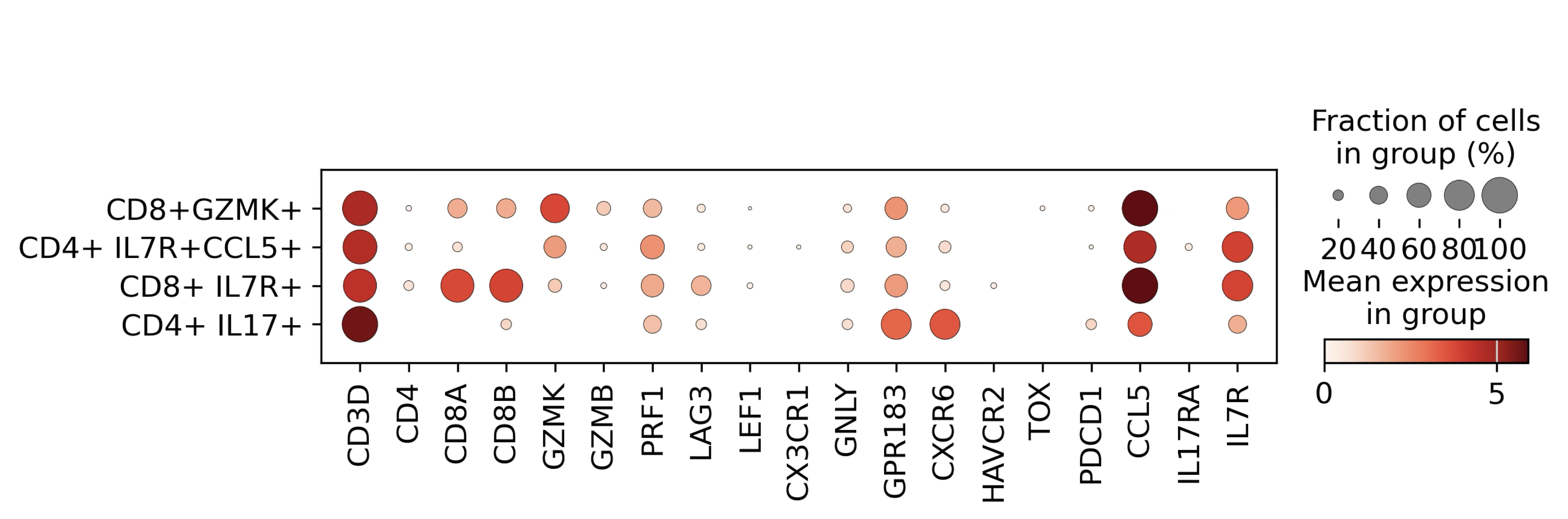

**Figure S18.** Marker gene expression of CD4-GNLY T cells. Related to **Fig 4.**

All CD4-GNLY T cells from the Zhang/Yu atlas, grouped by their predicted label. Cells annotated as CD8+ GZMK and CD8+ IL7R+ expressed CD8A and CD8B.

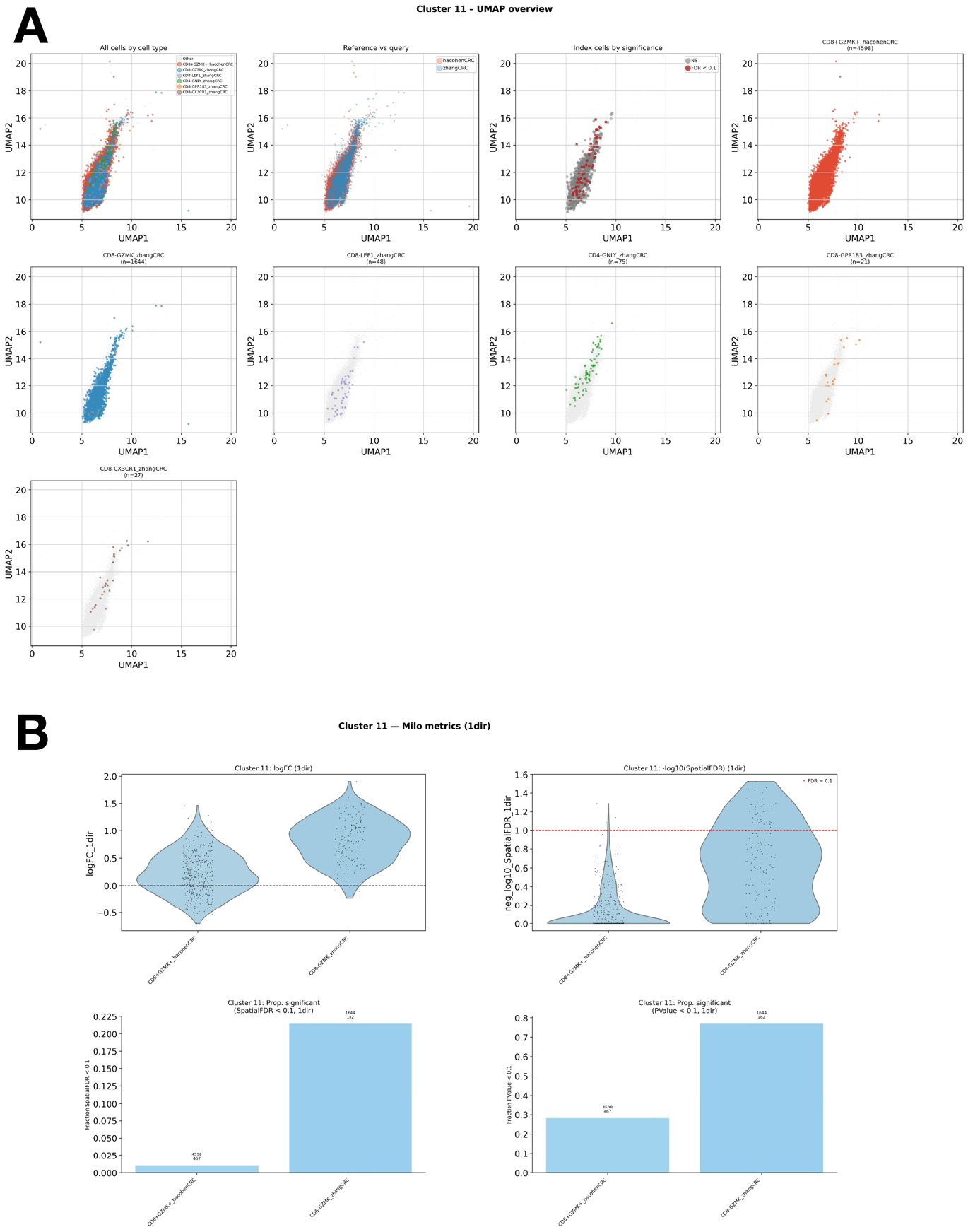

**Figure S19.** Inspection of Cluster 11 in CRC/CRC integration. Related to **Fig. 4.**

A) Cluster 11, colored by cell type and dataset.
B) DA testing results grouped by cell type within cluster 11.

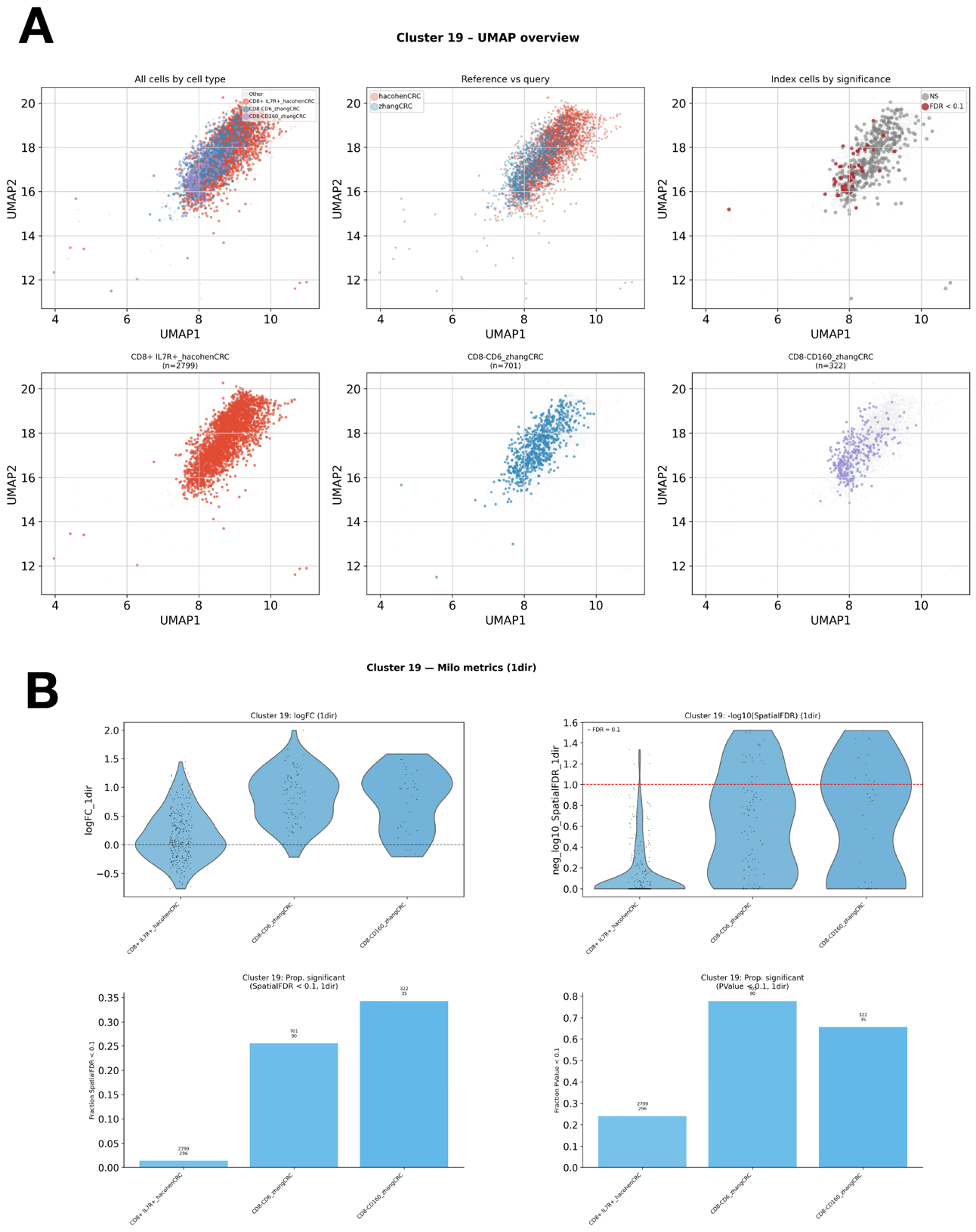

**Figure S20.** Inspection of Cluster 19 in CRC/CRC integration. Related to **Fig. 4.**

A) Cluster 19, colored by cell type and dataset.
B) DA testing results grouped by cell type within cluster 19.
